# The inflammasome adaptor *pycard* is essential for immunity against *Mycobacterium marinum* infection in adult zebrafish

**DOI:** 10.1101/2024.06.28.601189

**Authors:** Meri Uusi-Mäkelä, Sanna-Kaisa E. Harjula, Maiju Junno, Alina Sillanpää, Reetta Nätkin, Mirja T. Niskanen, Matti Nykter, Mika Rämet

## Abstract

Inflammasome regulates the host response to intracellular pathogens including mycobacteria. We have previously shown that the course of *Mycobacterium marinum* infection in adult zebrafish *(Danio rerio)* has similar features than the course of tuberculosis in human. In this study, we have investigated the role of the inflammasome adaptor *pycard* in *M. marinum* infection in zebrafish. We produced two zebrafish knock-out mutant lines for the *pycard* gene with CRISPR/Cas9 mutagenesis. While the zebrafish larvae devoid of *pycard* develop normally and have unaltered resistance against *M. marinum*, the loss of *pycard* led to impaired survival and increased bacterial burden in the adult zebrafish. Based on histological analysis, immune cell aggregates, granulomas, were larger in *pycard* deficient fish compared to wild type controls. Transcriptome analysis with RNA sequencing of a zebrafish haematopoietic tissue, kidney, suggests a role for *pycard* in neutrophil mediated defence as well as in haematopoiesis and myelopoiesis during infection. Transcriptome analysis of fluorescently labelled kidney neutrophils further supported the importance of *pycard* for neutrophil-mediated immunity against *M. marinum*. Genes associated with neutrophil degranulation, haematopoiesis and PI3K signalling were differentially expressed in the *pycard* deficient neutrophils when compared to wild type controls. All in all, our results indicate that *pycard* is essential for resistance against mycobacteria in adult zebrafish. Based on transcriptional profiling of *pycard* mutants, we postulate that *pycard* mutant phenotype is mediated in part via defects in neutrophil function including neutrophil degranulation.

**Author summary:** Inflammasome is a multiprotein complex which is a part of immune response especially against intracellular microbes. Tuberculosis, caused by *Mycobacterium tuberculosis*, is a global health problem causing 1.3 million deaths yearly. A natural fish pathogen, *Mycobacterium marinum*, causes a systemic infection in adult zebrafish resembling human tuberculosis including granuloma formation and an asymptomatic, latent state. Here, we have used zebrafish as a model to investigate the role of an inflammasome regulator *pycard* in resistance against *M. marinum* infection in zebrafish. We show that while *pycard* is dispensable for normal immunity against mycobacterial infection in larval stages, it has a central role in the immune defense against *M. marinum* in adult zebrafish. Based on our results, we suggest that this relates to blood cell formation, haematopoiesis and especially to the function of neutrophils. By analysing transcriptrional profile of *pycard* deficient neutrophils upon Mycobacterial challenge, it seems that *pycard* has a role in PI3K signalling and neutrophil degranulation. Our results indicate that *pycard* is essential for normal resistance against *M. Marinum* in adult zebrafish, and that it regulates neutrophil function during Mycobacterial infection.

## Introduction

Every year, approximately 10.6 million people are diagnosed with tuberculosis, and the disease causes 1.3 million deaths (1). Tuberculosis is caused by *Mycobacterium tuberculosis* (Mtb). Approximately 410 000 new tuberculosis cases are caused by either rifampicin resistant or multidrug resistant Mtb each year (1). Due to the limited efficiency of the Bacillus Calmette-Guérin (BCG) vaccine, new treatments and preventive methods are needed to eliminate tuberculosis (2). Most of the people infected with tuberculosis develop a latent infection, with dormant bacteria (3). In a latent infection, the mycobacteria can reside in granulomas without causing symptoms to the host, even for decades. It has been estimated that every fourth person is a carrier of the latent form of tuberculosis (1,4). The latent infection can reactivate to cause an active disease if the immune system is compromised (5).

Tuberculosis is transmitted by the intake of aerosolised bacteria from an infected person (6). Mtb travel to the lungs and are taken up by alveolar macrophages, where they survive by actively inhibiting the fusion of the phagosome with the lysosome (7). Preventing phagosome maturation is just one example of how Mtb manipulate the host’s protective pathways (8,9). Mtb is able to inhibit the activation of the inflammasome pathway, which is a key regulator of the post-translational activation of IL-1β and IL-18 (10,11). The inflammasome complex activates caspases which cleave the pro-inflammatory cytokines pro-IL-β, pro-IL-18 into their active form to promote the activation of host immune response (12). The inflammasome complex contains a receptor protein (either NLR (NOD-like Receptor) or AIM2 (Absent in Melanoma 2)), and an adaptor protein PYCARD (PYC and CARD domain containing, also known as ASC, Apoptosis-associated speck-like protein containing a CARD) which interacts with the interleukin activating caspases (12). Besides interleukin activation, inflammasomes can also activate Gasdermin D, which forms pores in the plasma membrane and can trigger a specialised form of cell death that results in the release of immune activators into the extracellular space, referred to as pyroptosis (13). Thus, inflammasomes serve as a frontline defence mechanism, and several components of the pathway have been shown to be essential for survival in mycobacterial infection (14,15).

Mtb has been shown to activate the NLRP3/ASC inflammasome through the ESAT-6 (early secretory antigenic 6 kDa (ESAT-6)) protein, which is essential for mycobacterium mediated phagosome maturation and mycobacterial virulence (16). Mtb causes damage to the plasma membrane, which induces the potassium efflux driven activation of the NLRP3 inflammasome in monocytes and macrophages (17). This can also result in Gasdermin D pore formation, pyroptotic cell death and release of the IL-1β into the extracellular space (17). Notably, the attenuated BCG *Mycobacterium bovis* fails to activate the NLRP3 inflammasome, likely due to the lack of the components of the region of difference 1 (RD-1) locus (18).

The inflammasome adaptor protein PYCARD connects the receptor protein and the caspase and thus is essential for inflammasome function. In addition, recent evidence shows that Pycard also has inflammasome independent roles in immunity, including chemokine regulation and T-helper cell polarisation (19,20). A number of cytokines are regulated by *PYCARD* via inflammasome dependent and independent mechanisms during *Porphyromonas gingivalis* infection in THP1 and U937 cell lines (20,21). Both the caspase and the Nlrp3 are dispensable for host survival in mouse models for tuberculosis, whereas Pycard knockout mice had impaired survival in an Mtb infection (15). The mechanism behind this decreased susceptibility remains unexplored. In some experiments, Pycard has been found to regulate the migration of adaptive immune cells in a Dock2 mediated manner (22). Uchiyama et al. (21) found that the induction of the Th17/Th1 cell response to *Listeria monocytogenes* infection was impaired in *Pycard^-/-^* mice.

The development of novel therapies relies on relevant model systems. The zebrafish (*Danio rerio*) offers a versatile platform to model the human immune system and disease genetics (24). The zebrafish shares over 70% genetic homology with human (25). Zebrafish also serve as a valuable model for tuberculosis research, as their natural pathogen, *Mycobacterium marinum*, is closely related to Mtb. The zebrafish/*M. marinum* model is been well established (26–31). Similar to humans, adult zebrafish infected with *M. marinum* develop an infection with chronic and latent phases with highly structured granulomas (32,33) Using this model, the importance of selected immune genes for resistance against mycobacterial infection can be studied (32,34–36). The adult zebrafish has both the innate and the adaptive arms of immunity, whereas larval zebrafish rely only on innate immune responses (37,38). Therefore, innate immunity can be studied using the zebrafish larvae, and, in turn, the interplay of the innate and adaptive immunities in the adult zebrafish. The zebrafish are also suitable for modelling human inflammasome activation (39–44).

Increasing interest has been directed towards the role of the inflammasome in adaptive immunity, as well as towards the inflammasome independent role of the adaptor protein Pycard. We have previously shown that *pycard* is upregulated in *M. marinum* infected adult zebrafish mutants with impaired immunity (45). To further study the role of the inflammasome in the defence against a mycobacterial infection, we generated two knockout *pycard* fish lines and studied their ability to defend themselves against a *M. marinum* infection. We found that adult zebrafish devoid of p*ycard* are more susceptible for mycobacterial infection compared to their wild type siblings. In addition, RNA-sequencing of infected adult zebrafish revealed novel genes and pathways relevant for *pycard*-mediated immune defence against a mycobacterial infection, including a number of transcription factors associated with haematopoiesis.

## Results

### Mutations generated with CRISPR-Cas9 lead to the elimination of the *pycard* transcript in zebrafish larvae

To study the role of inflammasome signalling in zebrafish, we generated two knockout mutant *pycard^-/-^* lines with the CRISPR-Cas9 technology (Supplementary figure 1A). The mosaic founder fish bearing mutations (AB background) were crossed with WT (TL background) fish, resulting in offspring that are heterozygous for mutations. The F1 heterozygotes carrying a mutation at the end of the first exon of the gene were selected to establish two stable knockout lines (Supplementary figures 1B, C and D). Both of the selected mutations result in a frameshift and a subsequent premature stop codon and termination of transcription (Supplementary figures 1C and D). In both cases, a predicted stop codon results in the deletion of at least the whole caspase recruitment domain (CARD, Supplementary figure 1). The expression of the *pycard* transcript was measured with quantitative PCR early in development in homozygous knockout and WT siblings. Both of the mutant lines had diminished *pycard* expression (Supplementary figure 1E and F). We considered these lines suitable for studying the loss-of-function phenotype of *pycard* in zebrafish.

### Loss of *pycard* does not impair immunity against *M. marinum* in larval zebrafish

Zebrafish larvae are protected solely by the innate immune system, as the T and B cells of the adaptive immunity develop only after around 20 days post fertilization (dpf) (37,38). To investigate whether *pycard* expression is required for resistance against mycobacterial infection in zebrafish larvae, we carried out yolk sack infections at 2-8 cell stage (Figure 1). A low dose of *M. marinum* infection did not result in compromised survival in the *pycard* larvae (Figure 1). As Pycard could be present in the oocytes from maternal transcripts, the infection experiment was repeated with larvae from homozygous (*pycard^-/-^*) and WT (*pycard^+/+^*) parents. In this setting, *pycard* was dispensable for immunity against low dose *M. marinum* infection (Figure 1). Thus, we conclude that *pycard* is not essential for the innate immune defence of larval zebrafish in a low dose mycobacterial infection. This supports earlier findings from Matty et al. (2019) (46), suggesting innate immune response against *M. marinum* infection in larval zebrafish is not dependent on Pycard.

**Figure 1.**
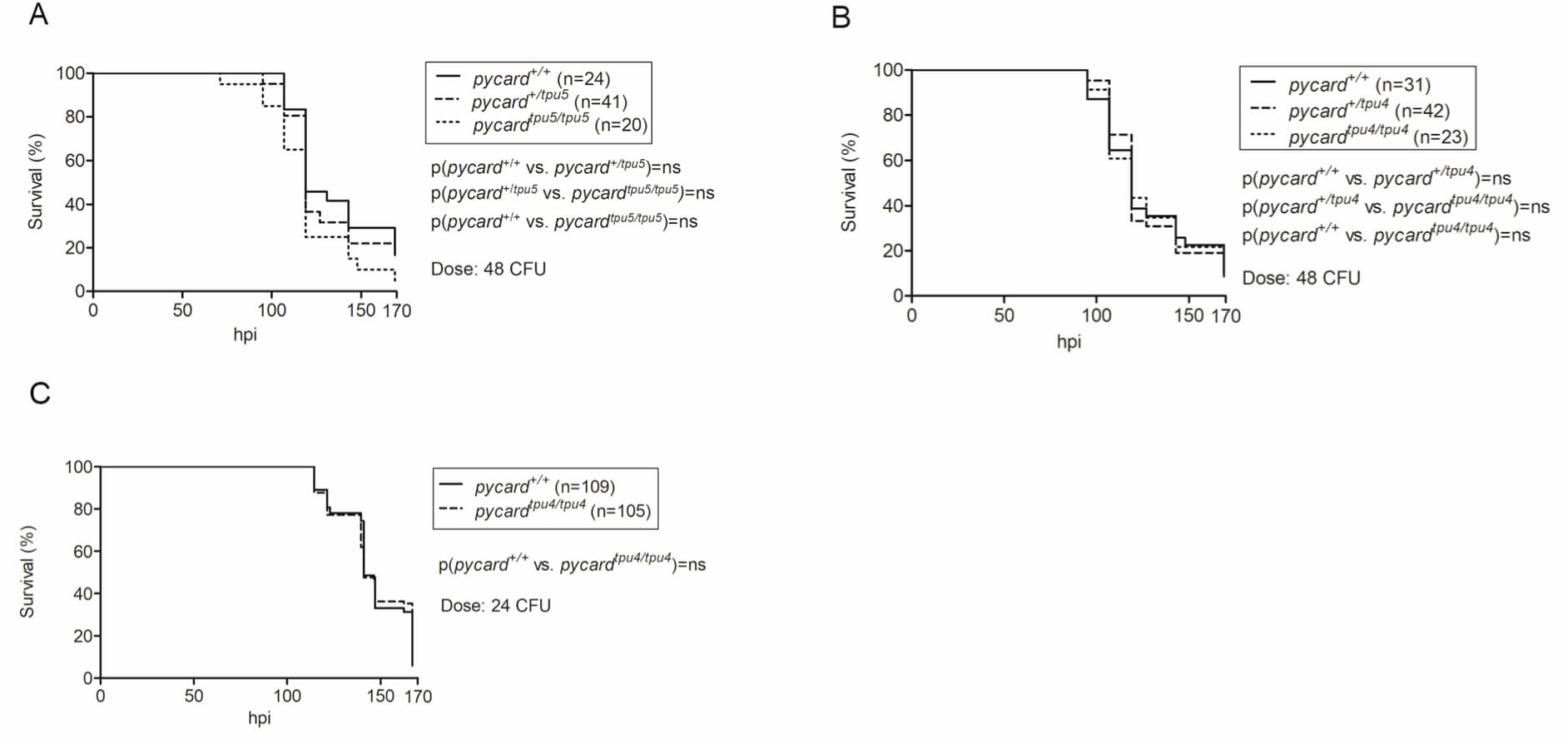
*pycard*^-/-^ larvae have normal survival in a mycobacterial infection. Embryos were infected with a low dose of *M. marinum* at the 2-8 cell stage and their survival was followed for 7 days. A) F2 generation results for the mutant line *pycard^tpu5^* (average dose 48 CFU, range 17-79 CFU). End point survival proportions *pycard^+/+^* 16.7%, *pycard^+/tpu5^*: 17,1%, *pycard^tpu5/tpu5^*: 5,0%) B) F2 generation results for the mutant line *pycard^tpu4^* (average dose 48 CFU, range 17-79 CFU). End point survival proportions *pycard^+/+^*: 9.7%, *pycard^+/tpu4^*: 11.9%, *pycard^tpu4/tpu4^*: 8.7%). C) Pooled results of three repeats for the experiment with the F3 generation of *pycard^tpu4^* (average dose 24 CFU, range 2-52 CFU). End point survival proportions *pycard^+/+^*: 5.8%, *pycard^tpu4/tpu4^* 12.4%). The average dose was measured from an infection solution plated on 7H10 agar plates. The survival data are presented as a Kaplan-Meier survival curve. The statistical analysis was done with the Log-rank test. hpi=hours post infection.

To determine whether *pycard* deficiency effects bacterial burden, we infected 2-1000 cell stage embryos originating from *pycard^+/tpu4^* F5 parents with a low dose of *M. marinum* (Figure 2). Bacterial copy number in 4 dpi (days post infection) larvae was similar in *pycard^tpu4/tpu4^*larvae compared to *pycard^+/tpu4^* and *pycard^+/+^* (median CFU 32805, 23812 and 32721, respectively (Figure 2). Thus, *pycard* expression is dispensable for the innate immune response against *M. marinum* in zebrafish larvae.

**Figure 2.**
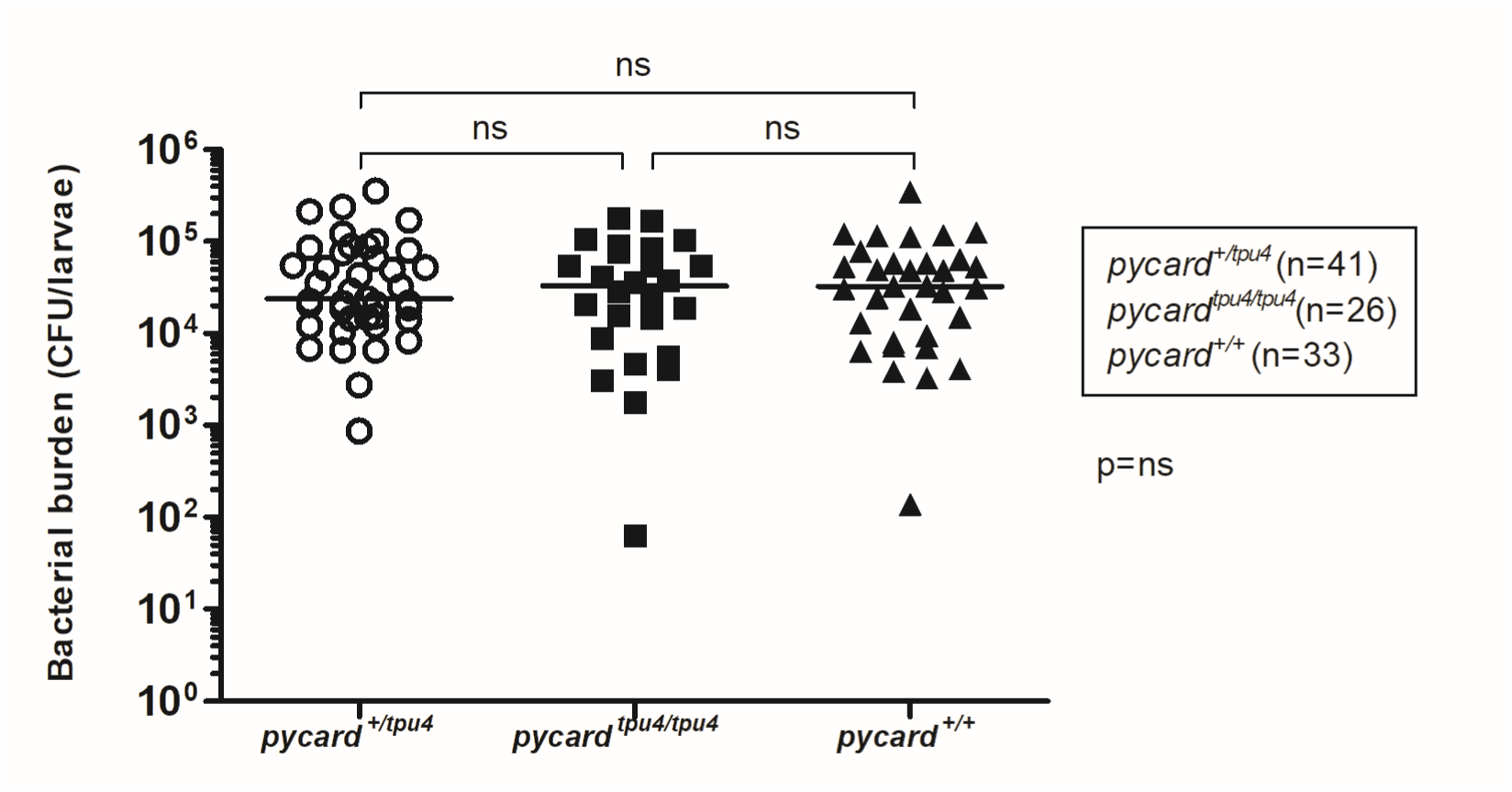
*pycard* expression does not affect bacterial burden in mycobacterial infection at 4 dpi in zebrafish larvae. Embryos originating from *pycard^+/tpu4^* F4 parents were injected with a low dose of *M. marinum* average dose (94 CFU, range 56-136 CFU) at the 2-1000 cell stage. Infected larvae were euthanized at 4 dpi, and DNA was extracted from the whole larvae. Larvae were genotyped and bacterial copy number was quantified using qPCR and *M. marinum* specific primers. The statistical significance of the results was analyzed with Kruskal-Wallis test. The line represents median.

### *pycard* is essential for the defence against an *M.* marinum infection in adult zebrafish

Inflammasomes are often regarded as a mechanism of the innate immunity. However, recent evidence points towards a role also in the adaptive immunity. Moreover, Nlrp3 and Pycard may have inflammasome independent roles (15,19,20,47). To this end, we studied whether *pycard* affects the immunity against *M. marinum* in adult zebrafish by following the survival during a low dose mycobacterial infection. The survival of the knockout *pycard^tpu5/tpu5^* fish was markedly impaired in a low dose infection with end point survival being 75.2% for *pycard^+/+^* and 40.6% for *pycard^tpu5/tpu5^* (p=0.002**) (Figure 3). To investigate whether reduced survival is attributed to compromised resistance or tolerance, we determined the bacterial burden of infected mutants and wild-type siblings. Adult zebrafish were infected with a low dose of *M. marinum* and bacterial burden of the fish was measured at 4 weeks post infection (wpi) with two independent mutant lines for *pycard* (Figure 3). We found that the bacterial burden was significantly increased in both mutants in internal organ block (for *pycard^tpu4^* medians were *pycard^+/+:^* 10045 CFU, *pycard^tpu4/tpu4^* 53426 CFU, p=0.0033**, for *pycard^tpu5^ pycard^+/+^*: 20349 CFU, *pycard^tpu5/tpu5^*: 48648 CFU, p=0.023*) (Figure 3). These data indicate that *pycard* expression is required for normal resistance against a low dose *M. marinum* infection in adult zebrafish. Results and bacterial doses from the individual experiments are shown in Supplementary figures 2 and 3.

**Figure 3.**
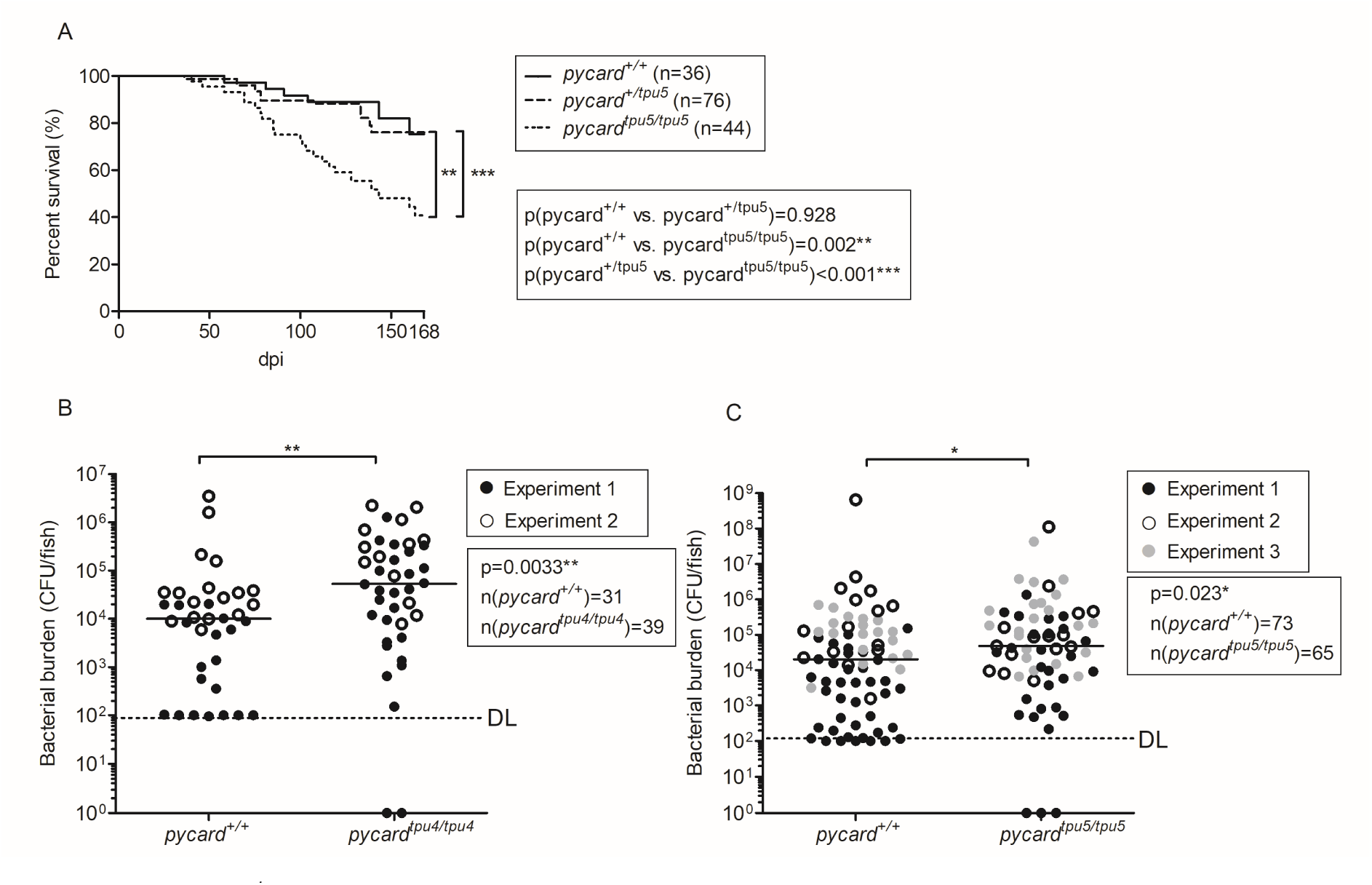
*pycard*^-/-^adult fish display reduced survival and a higher bacterial burden in a mycobacterial infection. A) Adult fish of the *pycard^tpu5^* line were infected with a low dose of *M. marinum* and their survival was followed daily (dpi = days post infection). The fish were genotyped post-mortem. The survival data are presented as a Kaplan-Meier survival curve. The statistical analysis was done with the Log rank test. The data have been pooled from two experiments. Individual experiments, with bacterial doses, are shown in Supplementary figure 2. Zebrafish from B) the mutant line *pycard^tpu4^* (Medians *pycard^+/+^*: 10045 CFU, *pycard^tpu4/tpu4^:* 53426 CFU, p=0.0033**) and C) the mutant line *pycard^tpu5^* (Medians *pycard^+/+^*: 20349 CFU, *pycard^tpu5/tpu5^*: 48648 CFU, p=0.023*). Fish were infected with a low dose of *M. marinum*, and the bacterial burden was analysed at 4 weeks post infection (wpi) from whole organ block DNA using qPCR with *M. marinum* genome specific primers. Individual experiments are presented with different symbols and are shown in separate graphs with the bacterial dose and the respective statistics in Supplementary figure 3. The line indicates the median. The data were analysed with the Mann-Whitney U-test, two tailed. For statistical purposes, samples in which the bacterial burden was below the detection limit (DL), *pycard*^+/+^ were designated a value of 100 CFU and for *pycard*^-/-^ a value of 0 CFU, respectively. Both sexes were included in the experiment in approximately equal numbers.

### *pycard^tpu4/tpu4^* knockout adult fish have a normal blood cell distribution

Previously, inflammasome components have been implicated in haematopoiesis and myelopoiesis (39,44). As *pycard* knockout fish had higher bacterial burden than their WT siblings (Figure 3), we investigated whether the altered resistance was associated with changes in the number of leukocytes. Thus, we collected the kidneys, which are the main site of haematopoiesis in fish, from both *M. marinum* challenged and mock (phosphate buffered saline (PBS)) injected adult zebrafish. At 4 wpi, the mutant fish presented similar number of precursor cells, monocytes and granulocytes and lymphocytes as WT siblings (Figure 4). Experiment was done twice, and pooled data are presented in Figure 4. As the number of cells remained similar to that in WT fish, it appeared that *pycard* does not affect leukocyte development in the adult zebrafish kidney. Individual experiments, including bacterial doses, and a full gating strategy are displayed in supplementary materials (Supplementary figure 4). Of note, this type of analysis cannot distinguish between different leukocyte sub-populations.

**Figure 4.**
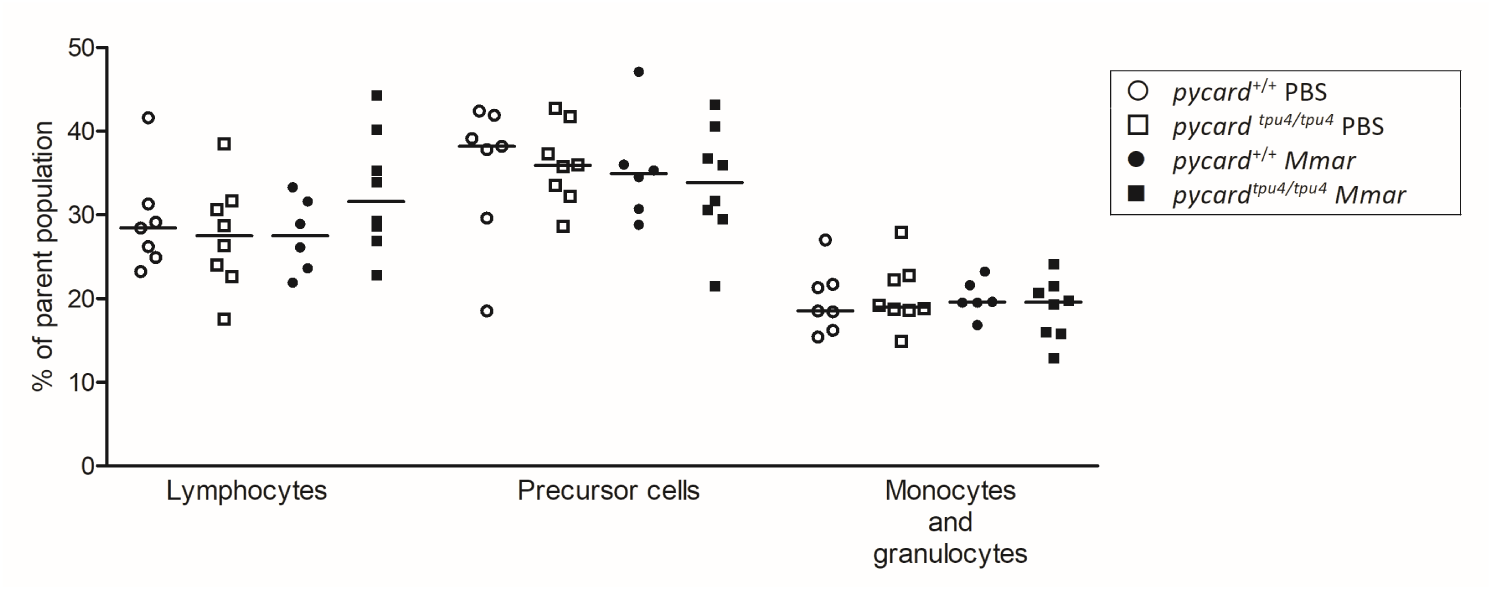
Unchallenged as well as *M. marinum* infected *pycard*^tpu4/tpu4^ mutants have normal distribution of blood cell populations. Adult zebrafish were either mock injected (PBS) or infected with a low dose (in experiment 1, mean dose 17 CFU, range 12-28 CFU, in experiment 2, mean 19 CFU, range 16-21 CFU) of *M. marinum* (*Mmar*). The whole kidney marrow was analysed at 4 wpi using flow cytometry. The results have been pooled from two experiments (n=3-5 in each group, per experiment). Each datapoint represents an individual fish. The line indicates the median. See supplementary figure 4 for the individual experiments including the bacterial dose and the gating strategy. Both sexes were included in the experiment in approximately equal numbers.

### *pycard* is widely expressed across tissues

To gain further information on the role of *pycard* in zebrafish with and without *M. marinum* challenge, the mRNA expression was analysed with qPCR from tissues and FACS sorted blood cell samples of AB WT fish. For blood cell analyses, fish were infected with a low dose of *M. marinum*, and samples were collected for FACS at 4 wpi. Highest median expression was seen in the spleen, gills, tailfin, gut, eyes, skin, muscle, and kidney (Figure 5A). The kidney and spleen are the main sites for haematopoiesis in zebrafish, whereas the gills, skin, tail, eyes, and gut are immunologically significant as they are exposed to the surrounding environment. In the granulocyte population which also harbours monocytes, *pycard* expression was decreased during the infection (Figure 5).

**Figure 5.**
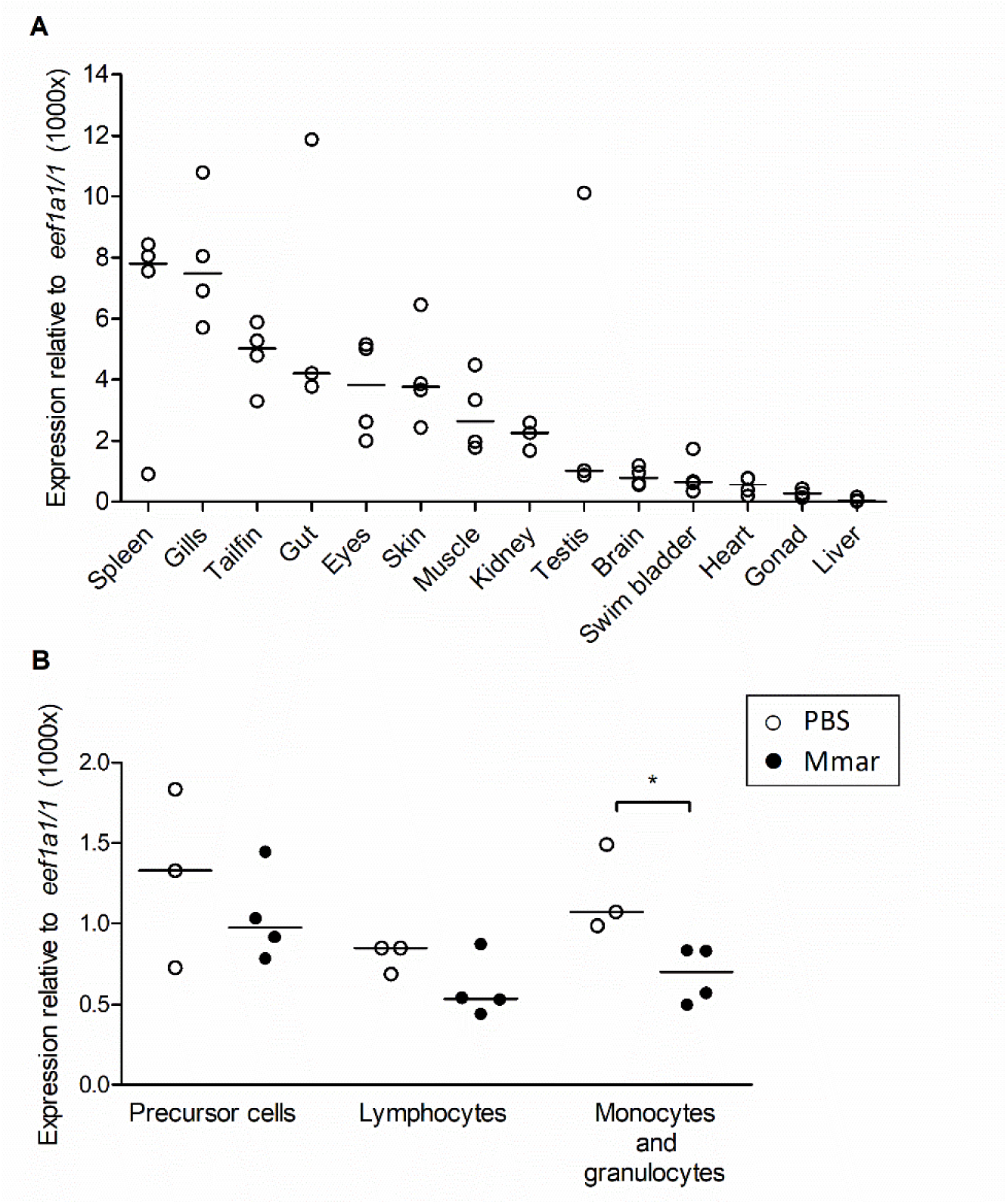
*pycard* is expressed in different tissues and blood cell types. A) Each datapoint represents the expression level of *pycard* in a single organ collected from a single AB fish (no treatment). B) AB Zebrafish infected with *M. marinum* (Mmar) (mean 28 CFU, range 21-34 CFU) display a decrease in *pycard* expression in their monocyte and granulocyte population. Fish were either infected with *Mmar* or mock injected with PBS. At 4 wpi, the fish were euthanised and their kidneys collected for FACS. RNA was extracted from the sorted populations and *pycard* expression was measured with qPCR and normalized to *eef1a1/1* expression. Each sample contained kidneys from three fish, pooled. Both male and female fish were used in both experiments. The line indicates the median. p(monocytes and granulocytes)=0.0286*, Mann-Whitney two-tailed)

### Granulomas in *pycard^tpu5/tpu5^* fish are larger than in WT siblings

Adult zebrafish develop mycobacterial granulomas with caseous necrosis, hypoxic core and a fibrous cuff, but fewer lymphocytes than mammalian granulomas (28,32,33). We next investigated whether lack of *pycard* expression affects granuloma formation in the adult zebrafish. The adult zebrafish were infected with a low dose (mean 64 CFU, range 53-76 CFU) of *M. marinum* and granulomas were characterised from the histological sections using Ziehl-Neelsen, Mallory’s trichrome or hypoxia staining from samples collected at 8 wpi. As shown in Figure 6, *pycard^tpu5/tpu5^*adult fish had larger granulomas compared to WT siblings (mean(WT)= 140.8 µm, mean(*pycard^tpu5/tpu5^*)=167.1 µm, p= 0.0217*). We also analysed the type of granulomas both in mutant and wild-type fish. Classification to nascent, necrotic and hypoxic granulomas was done as described in Myllymäki et al. 2018 (28). We also analysed whether granulomas had fibrotic capsule. There were not any notable differences in the distribution of different types of granulomas (Supplementary figure 6). These data suggest that *pycard^tpu5/tpu5^* fish are less capable of containing bacterial growth within the granulomas leading to increased bacterial burden and subsequently to a compromised immunity against *M. marinum*.

**Figure 6.**
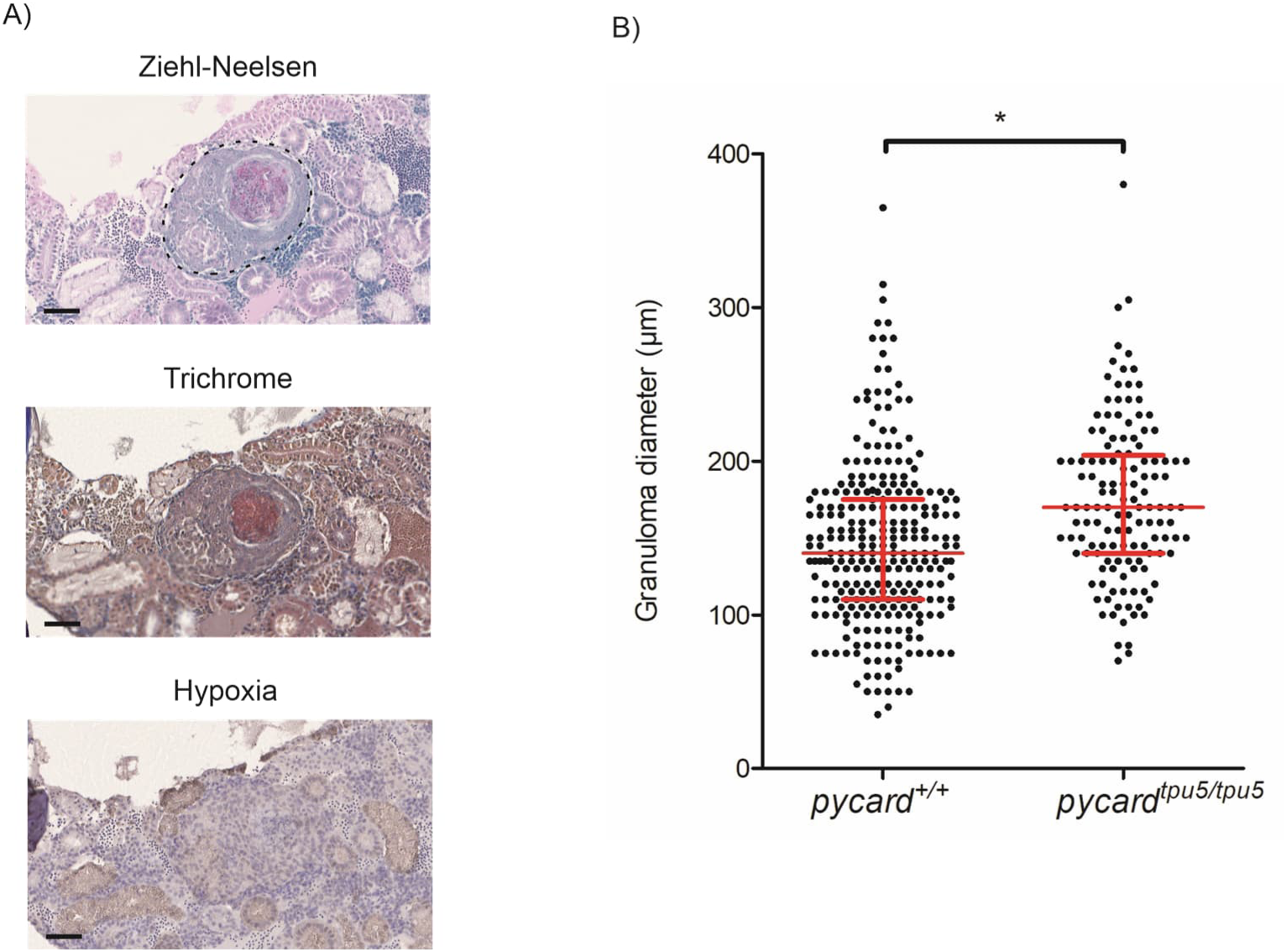
*pycard^tpu5/tpu5^* fish display an increase in granuloma size. Six WT and four *pycard^tpu5/tpu5^* fish were infected with a low dose (average dose 64 CFU, range 53-76 CFU) of *M. marinum*. At 8 wpi, the fish were processed for a histological analysis of granulomas with A) the Ziehl-Neelsen stain, Mallory’s trichrome stain and Hypoxia staining. The area which was considered to belong to the capsule of the granuloma is indicated in the Ziehl-Neelsen stain with a dashed line. The scale bar indicates 50 µm. B) The number and the characteristics of each granuloma were recorded for each fish. Using a linear mixed model, granulomas in *pycard^tpu5/tpu5^* were determined larger than in WT siblings (p=0.0217*) using R-package lme4, with fish as a random and genotype as a fixed factor. The line indicates the median and the interquartile range. Only male fish were used as female fish often present an increased number of small granulomas in the gonads, which complicates analyses. See also supplementary figure 5 for sizes of individual fish granulomas and for figure 6 for characterisation.

### Pycard affects the expression of haematopoietic transcription factors in an *M. marinum* infection

To identify the cause for the increased susceptibility of *pycard^-/-^*zebrafish to *M. marinum* infection, transcriptomes of *pycard^tpu4^*mutants as well as WT controls were analysed with RNA-seq from the kidney marrow at 4 wpi after a low dose of *M. marinum* (mean dose 31 CFU, range 16-48 CFU) or mock (PBS) injection. Based on RNA-seq, four genes were differentially expressed (DESeq2 (48,49), |log2 fold change| >1 between the groups, adjusted p value <0.05) without mycobacterial challenge, (Figure 7; Supplementary table 1). In turn, infected mutants showed 123 differentially expressed genes, in comparison to WT siblings, 21 of which were upregulated in the *pycard^tpu4^* mutant compared to WT and 102 downregulated (Figure 8; Supplementary tables 2-3).

**Figure 7.**
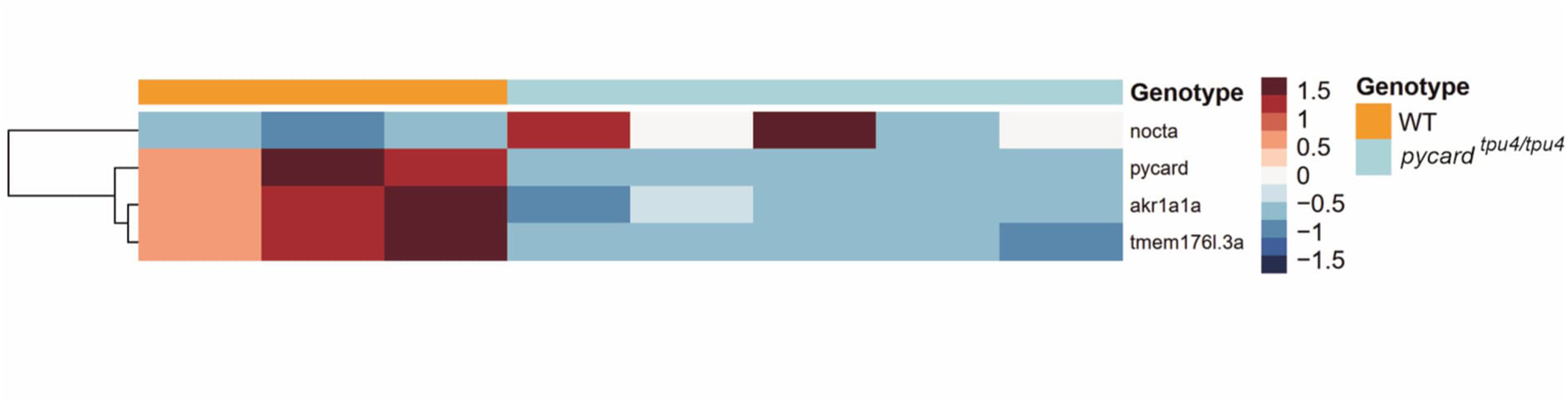
Heatmap of the RNA-seq results in *pycard^tpu4/tpu4^* adult zebrafish mock injected with PBS. Adult zebrafish from the *pycard^tpu4^* mutant line were mock injected, and at 4 wpi sacrificed for analysis. Whole kidney marrow was used for the RNA-seq analysis of male fish (n(WT)=3, n(*pycard^tpu4/tpu4^*)=5). See also Supplementary Table 1. Expressions are centered and scaled by row. Hierarchical clustering with Euclidean distance and complete linkage method was used to order genes.

**Figure 8.**
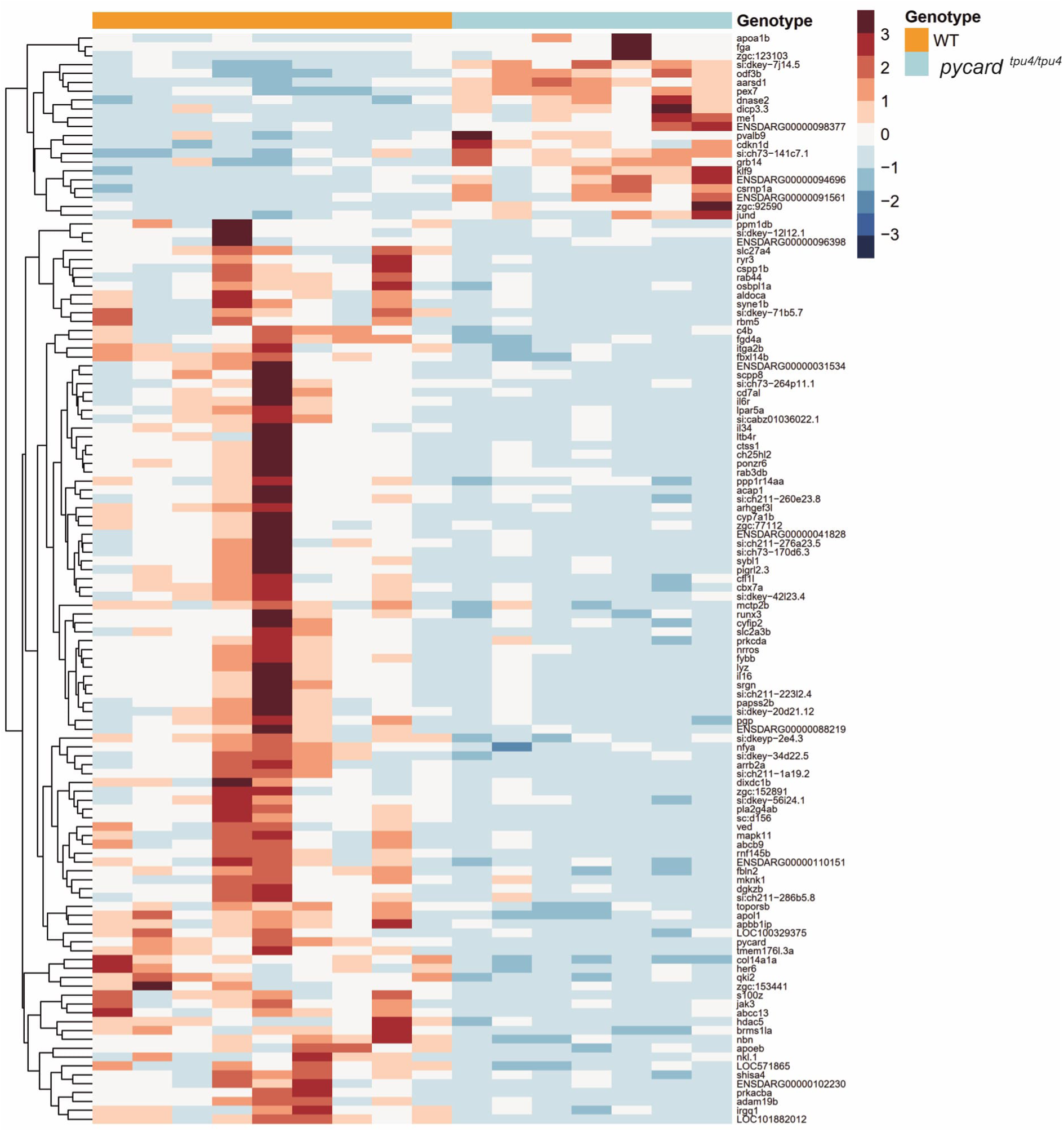
Heatmap of the RNA-seq results in adult *pycard^tpu4/tpu4^* zebrafish infected with a low dose of *M. marinum.* Adult zebrafish from the *pycard^tpu4^*mutant line were infected with a low dose of *M. marinum* (mean dose 31 CFU, range 16-48 CFU), and at 4 wpi they were sacrificed for analysis. Whole kidney marrow was used for the RNA-seq analysis in the male fish (n(WT)=9, n(*pycard^tpu4/tpu4^*)=7. See also Supplementary tables 2-3. Expressions are centered and scaled by row. Hierarchical clustering with Euclidean distance and complete linkage method was used to order genes.

We observed that a neutrophil marker, *lysozyme* (*lyz*) was downregulated in *pycard* mutants. In addition, other cell type specific genes, such as *cd7 antigen-like* (*cd7al*) (T-cell) (50), *Janus kinase 3 (a protein tyrosine kinase, leukocyte)* (*jak3*) (NK-cell), *integrin, alpha 2b* (*itga2b*) (thrombocyte) and *apolipoprotein Eb* (*apoeb*) (macrophages/myeloid cells) were among the downregulated genes (if not otherwise mentioned classified to cell types according to the scRNA-seq by Tang et al. 2017 (51), accessed via the online tool developed by Lareau et al. 2017 (52). These data suggest that *pycard* mutants may have compromised demand-adapted response related to these cell types.

Next, we looked whether the RNA-seq analysis gave indications for differences in immune-related gene expression. We observed that differentially expressed genes included several transcription factors, many of which are known to affect myelopoiesis in zebrafish, mouse or human. These include *ring finger protein 145b* (*rnf145b*) (53), *Kruppel-like factor 9* (*klf9*) (54), *chromobox homolog 7a* (*cbx7a*) (55), *nuclear transcription factor Y, alpha* (*nfya*) (56), *histone deacetylase 5* (*hdac5*) (57), *RUNX family transcription factor 3* (*runx3*) (58) and *cysteine-serine-rich nuclear protein 1a* (*csrnp1a*) (59). Based on the scRNA-seq data by Tang et al. 2017 (51), accessed via the online tool developed by Lareau et al. 2017 (52), majority of the downregulated genes (60 out of 102) in the *pycard^tpu4/tpu4^* fish are expressed in neutrophils (Supplementary table 2). The upregulated genes do not present such a definite pattern (Supplementary table 3). We also divided the differently expressed genes into functional categories (Supplementary tables 1-3).

After the RNA-seq analysis, we used the other mutant line, *pycard^tpu5^*, to validate the results. To this end, we performed either a mock injection (PBS) or a low dose infection (mean 24 CFU, range 16-32 CFU) to *pycard^tpu5/tpu5^*and WT siblings and analysed the gene expression from kidney tissue at 4 wpi for 10 selected genes with qPCR. The results for *pycard* itself, for a putative negative regulator of inflammasome, namely *transmembrane protein 176l.3a* (*tmem176l.3a*) (60), potentially a T cell activating gene *diverse immunoglobulin domain-containing protein 3.3* (*dicp3.3*), *malic enzyme 1* (*me1*) were replicated whereas the results for *aldo-keto reductase family 1, member A1a (aldehyde reductase)* (*akr1a1a*), *klf9*, *itga2b*, *lyz*, *nocturnin a* (*nocta*) and *cbx7a* did not differ between *pycard^tpu5^* mutants and controls (Supplementary figure 7) were not. *tmem176l.3a*, was differentially expressed after mock injection injection whereas *dicp3.3* and *me1* were differently expressed after *M. marinum* infection.

To further study the role of Pycard in the myelopoiesis we crossed homozygous *pycard^tpu4^* to transgenic zebrafish lines, *Tg(mpx:GFP)i114 (AB)* and *Tg(mpeg1.1:GFP)ka101 (AB)*, with fluorescent neutrophils and macrophages, respectively. Thus, we obtained zebrafish which are heterozygous for the mutation and carry fluorescent neutrophils or macrophages. We incrossed these fish to get homozygous, heterozygous and wild type larvae for *pycard^tpu4^*. At 3 dpf we wounded the tail fin in order to enhance haematopoiesis with a 30 gauge needle and imaged these fish to quantify the number of neutrophils and macrophages. In this setting, we saw no difference in the number of these cells between the wild type and the heterozygous or the homozygous mutants (Supplementary figure 8).

### The effect of Pycard on the transcriptomic profile of neutrophils during *M. marinum* infection

To study in more detail the effect of *pycard* deficiency on neutrophil in *M. marinum* infection, fish homozygous for *pycard^tpu4^*as well as WT controls, both having fluorescently marked neutrophils (*Tg(mpx:GFP)i114) (AB)* were infected with *M. marinum* (mean dose 95, range 69-118 CFU) and kidney marrow-derived neutrophils were sorted with FACS at 4 wpi (Supplementary figures 9A and B). There was no statistically significant difference in the relative neutrophil count between the WT controls and homozygous *pycard^tpu4^* fish (median 71.25 % vs. 63.95%; Supplementary figure 9B). Transcriptome of the sorted neutrophils was analysed with RNA-seq (Figure 9A). To verify the infection, whole organ blocks were collected, and bacterial burden was quantified (Supplementary figure 9C). As shown in Supplementary figure 9D, 20 most expressed genes in WT fish are dominantly expressed in neutrophils (according to the data by Tang et al. 2017 (51)). This indicates that FACS-based sorting of GFP expressing cells had yielded neutrophils.

**Figure 9.**
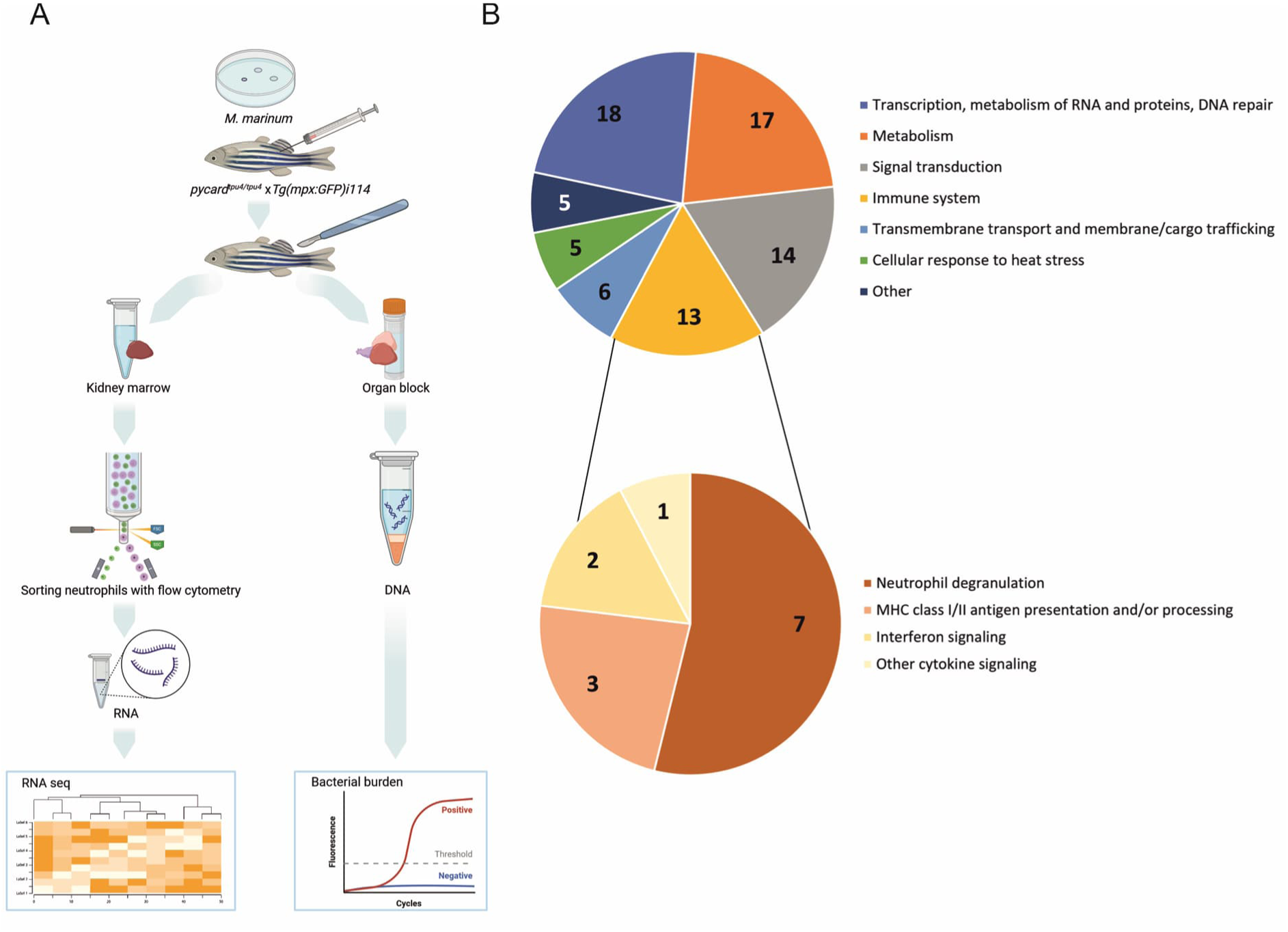
Transcriptomic analysis of kidney marrow-derived neutrophils and categorization of differentially expressed genes in adult *pycard^tpu4/tpu4^* zebrafish infected with *M. marinum*. A) Fish homozygous for *pycard^tpu4^* and WT controls crossed to *Tg(mpx:GFP)i114 (AB)* were infected with *M. marinum* (mean dose 95, range 69-118 CFU). At 4 wpi, fish were euthanized and dissected to obtain a kidney marrow and a whole organ block. From a suspended kidney marrow, fluorescent (GFP) neutrophils were sorted with flow cytometry and RNA was extracted for transcriptomic analysis with RNA-seq (n(WT)=3, (*pycard^tpu4/tpu4^*)=3. To monitor the effect of varying bacterial burden on transcriptomics, bacterial copy number was determined with qPCR from DNA extracted from the whole organ block. Illustration created with BioRender.com. B) All differentially expressed genes and immune system related genes were categorized according to Reactome (61). One gene can be assigned into several categories.

*pycard^tpu4/tpu4^* mutant zebrafish had 178 differentially expressed genes (DESeq2 (48,49) |log2 fold change| > 1 and the |fold change of medians| > 2 between the groups, p < 0.05 before adjustment) during *M. marinum* infection compared to WT their siblings (Supplementary table 4). Differentially expressed genes in *pycard^tpu4/tpu4^* mutants were categorized according to Reactome (61) (Supplementary table 5) (Figure 9B). 18 genes were related to transcription, metabolism of RNA and proteins or DNA repair, 17 to metabolism, 14 to signal transduction, 13 to immune system, 6 to transmembrane transport and membrane or cargo trafficking, and 5 to both cellular response to heat stress and to other functions. Of note, 4 genes, *platelet-derived growth factor receptor, beta polypeptide* (*pdgfrb*)*, regulatory associated protein of MTOR, complex 1* (*rptor*)*, Ras-related GTP binding Ca* (*rragca*) and *Scm polycomb group protein homolog 1* (*scmh1*) are on the Reactome pathway “PIP3 activates AKT signaling” (Supplementary Table 5). Genes related to immune system were categorized further (Figure 9B). Seven genes were related to neutrophil degranulation, three to MHC class I/II antigen presentation and/ or processing, two to interferon signaling and one to other cytokine signaling. The detailed list of the genes and their related functions can be found from Supplementary tables 4 and 5.

In *pycard^tpu4/tpu4^* mutants, 12 genes were significantly (adj. p-value <0.05) upregulated (Supplementary table 6) and 26 downregulated (Supplementary table 7). Among downregulated genes, 15 genes or their orthologs were related to immunity and 13 to PI3K-signaling (Supplementary table 7), whereas as 1 upregulated gene was associated to PI3K-signaling and 6 to immunity (Supplementary table 6). Based on the neutrophil RNA-seq data, the possible defect in *pycard^tpu4/tpu4^* neutrophils during *M. marinum* infection found in the kidney RNA-seq data may be related to PI3K/Akt signaling. Of note, PI3K/Akt signaling has been previously associated to Pycard and NLRP3 inflammasome (62–65).

## Discussion

In this study, we used CRISPR-Cas9 to produce two knockout zebrafish lines, devoid of the inflammasome adaptor gene *pycard*, and studied their phenotype during *M. marinum* infection. The qPCR analysis of the larvae showed that both of the *pycard* knockout lines (*pycard^tpu4^*and *pycard^tpu5^*) present diminished transcript levels (Supplementary figure 1). Moreover, infected *pycard^-/-^* larval zebrafish did not present altered survival or bacterial burden in *M. marinum* infection, indicating that *pycard* is dispensable for the innate immune response against *M. marinum* in zebrafish larvae (Figures 1 and 2). In turn, analyses of the adult zebrafish revealed an increased susceptibility in the mutated *pycard^-/-^*fish upon mycobacterial infection. In addition, the bacterial burden was increased in *pycard* mutant lines (*pycard^tpu4^* and *pycard^tpu5^*) (Figure 3). Altered resistance was accompanied with changes in transcription especially in the genes related to neutrophil function (Figure 8, Supplementary table 3). A previous zebrafish study (46) showed that the survival or bacterial burden of *pycard* knockout larvae was not markedly affected in an *M. marinum* infection. Our data indicate that the role of *pycard* is more important in mycobacterial immunity in adult zebrafish compared to larvae. This may be due to differences in the immune response but also due to very different infection kinetics between these two models. However, we consider the adult model informative as it includes both the innate and adaptive immune responses and therefore reflects better mammalian models of tuberculosis.

The inflammasome adaptor protein coding gene *pycard* is required for inflammasome signalling and IL-1β production in zebrafish (66). In mouse, the lack of Pycard affects the course of mycobacterial infection (14,15). In addition, PYCARD has been suggested to regulate the immune response independent of the inflammasome (19,21,47,67). In a mouse model, Pycard deficient mice have reduced survival against Mtb H37Rv infection (14,15). There were some differences in the phenotypes between these studies, however. Whereas McElvania Tekippe et al. (2010) found that survival from Mtb H37Rv strain infection was dependent on Pycard but not Caspase1, Mayer-Barber et al. (2010) observed a decreased survival in Caspase1 deficient mice (14,15). There is no obvious explanation for this discrepancy although this could be a result of different infection procedure used in the two studies (Mayer-Barber et al. 50-100 CFU, McElvania Tekippe et al. 250-350 CFU) (14,15). In addition, McElvania Tekippe et al. (2010) found that in Pycard deficient mice, a number of bacteria are found outside the granulomas due to an inability to contain the mycobacteria in the granulomas (15). In our study, *pycard*^tpu5/tpu5^ adult zebrafish presented larger granulomas than WT siblings (Figure 6). This could be due to for example the granulomas being less able to contain the bacteria, leading to infection progressing faster, which would be in line with higher bacterial burden as well as earlier death in the survival of the adult fish.

Based on our RNAseq analysis and further independent validation with qPCR, there were only few genes whose expression was affected by the loss of *pycard* in the kidney marrow of unchallenged adult zebrafish. In addition to *pycard* itself, the expression of *tmem176l.3a* was downregulated. Of note, the mouse ortholog Tmem176b has been shown to be a negative regulator of inflammasome signalling (60). *Tmem176l.3a* was also downregulated after *M. marinum* infection in the kidneys and neutrophils of the *pycard* deficient fish.

During *M. marinum* infection, over half of the (60 out of 102) downregulated genes in our data of the differentially expressed genes are expressed in neutrophils according to the scRNA-seq data by Tang et al. (2017) (51) (Supplementary table 3). Neutrophils are known to play a dual role in tuberculosis in mouse models (68,69). As neutrophils are the most numerous immune cell type in blood, they are often among the first responders to an infection and thus shape the initial response by initiating cytokine signalling. However, in an active infection, prolonged neutrophil activation can cause damage in host tissues (reviewed by Kroon et al. 2018) (57). Using zebrafish larvae, Kenyon et al. (2017) found many inflammasome-related genes upregulated in neutrophils at the early stage of an *M. marinum* infection in zebrafish larvae (71).

Inflammasome has been indicated in regulating haematopoiesis in zebrafish, mice and humans (39,44,72). Based on studies employing morpholino silencing of *pycard* in zebrafish larvae, it was suggested that the inflammasome cleaves the main haematopoietic transcription factor GATA binding protein 1a (Gata1a) into an active state (44). According to their study, Gata1a, in combination with Spi-1 proto-oncogene b (Spi1b), is responsible for the decreased numbers of macrophages and neutrophils in zebrafish larvae in a demand-driven haematopoiesis in morphant zebrafish (44). Frame et al. (2020) showed in a similar zebrafish model for *pycard* silencing, that inflammasome stimulation increased multilineage hematopoietic colony-forming units and T cell progenitors (39). Frame et al. (2020) also showed an increase in *interleukin 6* (*il6*) expression after exogenous *interleukin 1, beta* (*il1b*) induction by glucose, a signal which has been associated with promoting haematopoietic stem cell differentiation (39,72). Supporting this, our kidney RNA-seq data show downregulation of il6 receptor (*il6r*) in *pycard^tpu4/tpu4^* mutants after an *M. marinum* challenge.

There were also other signs suggesting compromised haematopoiesis upon mycobacterial challenge in *pycard^tpu4/tpu4^* mutants. These include reduced expression of the myelopoiesis and haematopoiesis associated transcription factors *rnf145b* (53), *nfya* (56), *hdac5* (57) and *runx3* (58), and elevated expression of *csrnp1a* (59). However, we are unable to determine in which specific cell types these transcription factors are expressed in our kidney samples. Also, our transcriptome analysis of the *pycard* deficient neutrophils shows significant downregulation of several (9 out of 26) haematopoiesis-associated genes. PI3K/Akt signaling associated genes and processes were enriched in our neutrophil-derived RNA-seq data. Among such enriched processes are haematopoiesis, neutrophil degranulation and cell migration (73,74). It has been shown in zebrafish that *pten*, a PI3K regulator (74), has a role in proliferation and differentiation of haematopoietic stem cells (75).

Tyrkalska et al. (44) showed decreased number of neutrophils and macrophages in unchallenged *pycard (asc)* morphants. However, according to Lozano-Gil et al. (2022) myeloid inflammasome does not affect haematopoiesis (76). In our setting, we could not see differences in neutrophil or macrophage counts in *pycard* mutant larvae post wounding, perhaps due to maternal RNA or other compensatory mechanisms. Neither was the presence of impaired haematopoiesis observed in the neutrophil counts determined with FACS from infected fish. This effect can be dependent on the progression of infection and would require further analysis. However, these results are supported by the fact that the neutrophil marker *mpx* used as a marker in our experiments did not show decreased expression due to *pycard* deficiency in our kidney or in our neutrophil RNA-seq data.

Several genes which were differentially expressed in infected *pycard^tpu4/tpu4^* compared to WT controls have been associated with mycobacterial infection. Oxidative stress and reactive oxygen species (ROS) are known to be critical for host defence in tuberculosis, and ROS has also been suggested to regulate inflammasome activation in mice in acute lung injury model (77,78). Genes associated with ROS-formation were malic enzyme 1, (*me1*) which was also confirmed with qPCR with the other mutant line *pycard^tpu4/tpu4^,* and negative regulator of ROS (*nrros*) (79,80). We consider downregulation of *nrros* as a compensatory effect for compromised resistance. *interleukin 34* (*il34*), was also downregulated in our kidney RNA-seq data and is essential in macrophage migration in zebrafish (81). Moreover, IL34 stimulated macrophages were shown to be more resistant to mycobacterial entry and more efficient in phagolysosomal trafficking of *M. marinum* in *Xenopus laevis* (82). The zebrafish leukotriene B4 activation mediates increased susceptibility to mycobacterial infection, and the ltb4 inactivating enzyme ltb4dh counteracts this increase (83,84). In the same study, inhibiting the ltb4 receptor rescued the susceptible phenotype of ltb4dh deficient zebrafish (84). *leukotriene B4 receptor* (*ltb4r*) was downregulated in the *pycard^tpu4/tpu4^* mutants in our RNA-seq. In addition, we observed significant downregulation of NK-lysin 1 (*nkl.1*). *nkl.1* belongs to a family of proteins which have been indicated in the elimination of *M. tuberculosis*, although the direct homology of the *nkl.1* and human granulysin remains to be shown (85). In addition, we observed notable dysregulation of one gene associated with platelet activation. We observed an increase in the expression level of fibrinogen alpha (*fga*). Recently, Hortle et al. (86) showed, that platelet activation during infection compromises the host immunity in a mycobacterial infection in larval zebrafish. In their study, *fga* knockout larval zebrafish fish presented with a decreased bacterial burden (86). Correspondingly *fga* was upregulated in our data. Platelets are known to be important for the immunopathology of tuberculosis, as they affect other immune cells and especially monocytes, to increase activation and enhance phagocytosis (87,88). These data suggest that *pycard^-/-^*zebrafish present a number of changes in their transcriptome indicating a compromised host protective immune response in mycobacterial infection.

According to transcriptomic analysis of *pycard^tpu4/tpu4^*neutrophils, the most prominent number of immune system related differentially expressed genes were related to neutrophil degranulation, an essential mechanism in the defence against pathogens. Pathogen induced inhibition of degranulation is associated to more severe outcome, whereas excess degranulation can damage host tissue. (89) We have previously shown the induction of neutrophil degranulation associated genes in *M. marinum* infection in adult zebrafish (45). In patients with active tuberculosis, genes related to neutrophil degranulation were downregulated compared to latent infection, referring to an inhibition of degranulation (90). In our current analysis, PI3K-signaling-related genes were differentially expressed in *pycard^tpu4/tpu4^* neutrophils. PI3K-signaling is known to regulate neutrophil activation (73), including degranulation (89).

Our results propose a role for *pycard* in defence against mycobacterial infection through regulation of a number of haematopoiesis and myelopoiesis associated genes and genes associated with neutrophil defence. Whether the dysregulated genes directly or indirectly associate with *pycard* remains to be studied.

We show that *pycard* expression is required for normal immunity against *M. marinum* in zebrafish. Our data highlight the difference between the larval and adult zebrafish models. We show that *pycard* mutants form granulomas in a similar manner to WT control siblings. However, the granulomas of the *pycard^tpu5/tpu5^*fish are larger suggesting that *pycard* mutants have compromised ability to restrict bacterial growth in granulomas. There are several transcriptional changes in mutants linking *pycard* function to demand-driven haematopoiesis and PI3K/Akt signaling. We postulate that *pycard* mutant phenotype is mediated in part via defects in neutrophil function like in neutrophil degranulation.

## Materials and methods

### Resource availability

Further information and requests for resources and reagents should be directed to the lead contact Mika Rämet.

### Materials availability

The CRISPR-Cas9 zebrafish lines *pycard^tpu4^ and pycard^tpu5^* generated in this study will be made available on request, but we may require a payment and/or a completed Materials Transfer Agreement if there is potential for commercial application.

### Data and code availability

The RNA-seq datasets from the adult zebrafish kidney (PBS injected controls and *M. marinum* infected samples) and from the adult zebrafish kidney neutrophils (*M. marinum* infected samples) have been submitted to the NCBI Gene expression omnibus (91,92) under the GEO Series accession numbers GSE189627 (https://www.ncbi.nlm.nih.gov/geo/query/acc.cgi?acc=GSE189627) and GSE270136 (https://www.ncbi.nlm.nih.gov/geo/query/acc.cgi?acc=GSE270136).

### Zebrafish lines and maintenance

All maintenance and experiments were done in accordance with the Finnish act on the protection of animals used for scientific or educational purposes (497/2013) and the EU Directive on the protection of animals used for scientific purposes (2010/63/EU). Permits for experiments were applied from the Regional State administrative agency (ESAVI/10079/04.10.06/2015 ESAVI/11144/04.10.07/2017, ESAVI/2776 /2019 and ESAVI/7251/2021). Before infection experiments, adult fish were maintained in a flow through system (Aquatic Habitats, Florida, USA). Fish were fed once a day with suitable granularity of Gemma Micro feed (Planktovie, Marseille, France). The light/dark cycle was 14/10 h in all laboratories and incubators. Embryos were kept in embryonic medium (5 mM NaCl, 0.17 mM KCl, 0.33 mM CaCl2, 0.33 mM MgSO4, 0.00001% Methylene Blue) in a 28.5°C incubator until 6 days post fertilization (dpf) and fed starting from 5 dpf.

Infected adult fish were kept in a separate laboratory in another flow through system (Aqua Schwartz mbH, Gönningen, Germany). Infected embryos were kept in 24-well plates in individual wells to prevent the spread of the infection. During infections, the wellbeing of the fish was followed at least once a day and fish exhibiting symptoms, signs of pain or discomfort were euthanised with an overdose of a Tricaine anaesthetic (Ethyl 3-aminobenzoate methanesulfonate, (Merck, Kenilworth, New Jersey, USA)). Adult fish used for experiments were 3 to 10 months old. Fish lines used were AB wild type (WT) or CRISPR-Cas9 mutants generated in house. CRISPR-Cas9 mutated fish were outcrossed to TL WT (*Tüpfel long fin, gja5b^t1/t1^, lof^dt2/dt2^*) so their background was AB x TL. All fish used in the experiments were the offspring of heterozygous parents, except the larvae used in the F3 survival experiment, as indicated in the text. The lines were assigned the Zebrafish information network (zfin.org) identifiers *pycard^tpu4^* and *pycard^tpu5^*. Each result indicates which line has been used for the experiment. The transgenic zebrafish lines, *Tg(mpx:GFP)i114 (AB)* (93) and *Tg(mpeg1.1:GFP)ka101 (AB)*, used to produce *pycard* mutants with fluorescent neutrophils or macrophages, respectively, were obtained from the European Zebrafish Resource Center (EZRC, Karlsruhe Institute of Technology, Eggenstein-Leopoldshafen, Germany).

### CRISPR-Cas9 mutagenesis and genotyping

The design procedure of our in house produced CRISPR-Cas9 mutant lines has been described in Ojanen et al. 2019 (36). Briefly, several online tools were used for selecting several guide RNA (gRNA) targets for *pycard* (ENSDARG00000040076.8). The gRNA was synthesised with the MEGAShortScript T7 transcription kit (Thermo Fischer Scientific, MA, USA). The in house produced Cas9 protein was obtained from the Tampere Facility for protein services. Injections were performed with borosilicate capillary needles generated with Flaming/Brown micropipette puller (Sutter, Novato, CA, USA), using a micro injector (PV830 Pneumatic PicoPump, World Precision Instruments, Sarasota, Florida, USA). 130 pg of gRNA and 250 pg of the Cas9 protein were co-injected into a 1-cell stage fertilized embryos of AB WT fish with phenol red (pn. 114537-5G, Merck) as a tracer dye. The CRISPR-injected F0 generation was crossed with TL WT fish, and from this F1 generation founder fish carrying frameshift causing mutations were identified with a heteroduplex mobility assay and then by Sanger sequencing. After identification of the desirable mutations, the F1 heterozygous mutant fish were incrossed to get the generation F2 (25% WT, 50% Heterozygous, 25% Homozygous mutants). F3 and F4 generations were used for larval experiments where the effect of the functional maternal *pycard* mRNA could be excluded. In bacterial burden analysis of larvae, F6 generation was used.

Fish used for experiments were genotyped by assessing the restriction fragment length with agarose gel electrophoresis, after digestion of a PCR amplicon with the CseI restriction enzyme (ThermoFisher Scientific, Waltham, Massachusetts, USA). DNA extractions for genotyping were always done either from whole larvae, or from adult tailfins excised under anaesthesia. Briefly, samples were lysed with a standard lysis buffer (10 mM Tris (pH 8.2), 10 mM EDTA, 200 mM NaCl, 0.5% SDS) with Proteinase K (ThermoFisher Scientific) (0.2 mg/ml) for a minimum of 2h in a 55°C water bath. For larval bacterial burden analysis, fixed lysis time of 21,5 hours was used. Two volumes of absolute ethanol were added, and DNA was precipitated at -20°C for minimum of 30 minutes. The DNA was then pelleted for 20 minutes at 16000 x g in a microcentrifuge. The pellet was washed with 200 µl of 70% ethanol before suspending it in nuclease free water. PCR was done with the Dream Taq Hot Start polymerase (ThermoFisher Scientific) with gene specific primers (F: 5’-GACCCAACTGTGAGGAACCATG-3’, R 5’-GCTTTCTTCAGACTTAAACGCCTTC-‘3). Digested fragments were separated on a 2% (w/v) agarose gel electrophoresis.

### Adult fish experiments

Offspring of a heterozygous fish crossing were used for the survival and bacterial burden assays and the fish were injected without knowing the genotypes beforehand, with the exception of fish injected for RNA sequencing from neutrophils. Fish were infected with a low dose of *M. marinum* intraperitoneally, under 0.02% Tricaine anaesthesia. The well-being of the fish was followed minimum once per day. Fish exhibiting symptoms of mycobacterial infection, signs of pain (as per humane end point) or discomfort were euthanised and collected for genotyping analysis. Symptoms included skin lesions, upturned scales, changes in swimming or behaviour, gasping for breath or lack of responsiveness.

### *M. marinum* infections

*M. marinum* (ATCC 927 -strain) was used for all infections. The bacterial preparation and injection procedure have been described previously (32). Briefly, the bacteria were inoculated from BD Difco Middlebrook 7H10 plates (BD Biosciences, NJ, USA) into 7H9 media (BD Biosciences) and grown at 29°C protected from light. After 72 hours, the culture was passaged to OD_600_=0.07 and allowed to reach the logarithmic growth phase. For larval infections, bacteria were pelleted and resuspended in a desired volume of 2% polyvinylpyrrolidone in phosphate buffered saline (PBS). Phenol red was used as a dye to visualize the injection, and a 1 nl injection volume was calibrated using a halocarbon oil droplet on a microscope scale bar. For survival analysis, larval zebrafish were injected with *M. marinum* or a PBS mock injection mix into the yolk sack at the 2-8 cell stages, and dechorionated at 1 dpf. For bacterial burden assay, embryos were injected with *M. marinum* at the 2-1000 cell stage. For adult fish, bacteria were pelleted and resuspended in PBS with 10% phenol red. Anaesthetised adult fish were injected with 5 µl of injection solution intraperitoneally using a 30 G needle. Both with larval and adult zebrafish, injection doses were plated onto 7H10 plates (BD Biosciences) during the procedure to verify the dose.

### RNA-extractions

A gene expression analysis of adult zebrafish tissues and larval zebrafish was done with quantitative PCR (qPCR). For the organ collection, adult AB zebrafish were euthanised and their organs collected with tweezers into PBS containing microcentrifuge tubes. Larval zebrafish were homogenized by pipetting. Zebrafish organs and larval zebrafish were homogenized by pipetting or with a needle, and a syringe. For RNA sequencing from a kidney marrow and qPCR, kidneys were homogenized in tubes containing an RNA preserving buffer, 6 ceramic beads (2.8mm ∅, Omni International, GA, USA), 6.5m/s for 2 x 30s on dry ice with FastPrep-24™ 5G bead beating grinder and lysis system (MP Biomedicals LCC, CA, USA). For RNA sequencing from neutrophils, kidney marrow was homogenized by pipetting prior to cell sorting and cell suspensions were homogenized using a syringe and a needle. RNA extractions were done with RNeasy Mini Plus or Micro Plus kits (Qiagen, Hilden, Germany). For the RNA expression experiments in the organs of the adult zebrafish, DNA removal columns were not used, instead the contaminating genomic DNA was removed with Rapid Out DNA removal kit (ThermoFisher Scientific).

### quantitative PCR analyses

For qPCR, cDNA was synthesised with a Sensifast cDNA synthesis kit (Bioline, Meridian bioscience, OH, USA). Transcript quantitation was performed with the PowerUp Sybr Green master mix (ThermoFisher Scientific) with a BioRad CFX96 Real-time PCR detection system (BioRad). Primers for qPCR are indicated in Table 1. Cycle threshold (Ct) values were normalized to the Ct values of the *eef1a1/1* transcript (94)), and amplicon sizes were confirmed with a gel run. The program cycling parameters were 50°C 02:00 (mm:ss), 95°C 02:00, 40 cycles of 95°C 00:03 and 59-60°C 00:30, followed by melt curve from 55°C to 95°C at 0.5°C increment. All primers are listed in table 1. All samples were measured in duplicate.

**Table 1.**
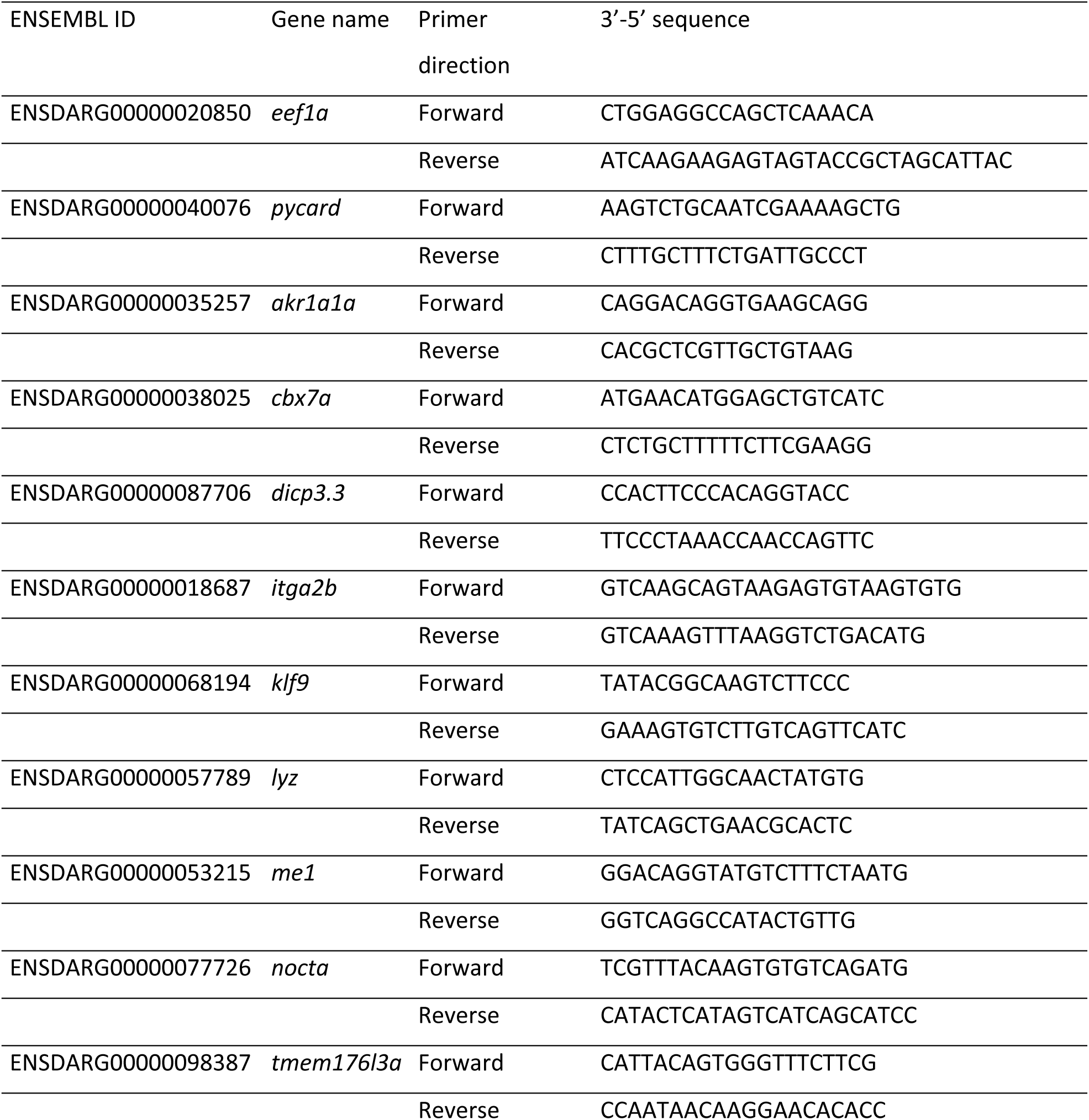
qPCR primers used in the study

### The determination of the bacterial burden

At 4 wpi, fish were euthanised with an overdose of Tricaine. Fish were pinned on a polystyrene piece and the whole organ block was released with tweezers and a spatula. The organ block was collected into a screw cap tube with 6 ceramic beads. Samples were homogenized in tubes containing the Tri-reagent (MRC Inc. OH, USA) 6.5 m/s for 2 x 30 s. on dry ice with a FastPrep-24™ 5G bead beating grinder and lysis system. DNA was extracted as in Parikka et al. *M. marinum* genome copies were quantitated with genome specific primers (primers F: 5’-CACCACGAGAAACACTCCAA-3’, R: 5’-ACATCCCGAAACCAACAGAG-3’ (32)) against a standard curve (1:5 dilution series) of a sample of known bacterial burden. The measurement was done with a Sensifast Sybr no-ROX kit (Bioline) in a Bio-Rad CFX96 Real-time PCR detection system, with up to 1 µg of DNA as template. The program used was 95°C 05:00, followed by 40 repeated cycles of 95°C 00:05, 65°C 00:10, 72°C 00:10, followed by a melting curve from 55°C to 95°C at 0.5°C increment. The correct size of the amplification product was confirmed with agarose gel electrophoresis. To determine the bacterial burden in larval zebrafish, larvae were euthanized at 4 dpi. After extracting DNA and genotyping larvae according to a protocol described for CRISPR-Cas9 mutagenesis and genotyping, number of *M. marinum* copies was quantified as described above.

### Flow cytometry and cell sorting experiments

Adult zebrafish were euthanised with an overdose of Tricaine anaesthetic. The fish was placed on a piece of polystyrene, cut with a scalpel, and pinned open. To determine the blood cell population, the kidney was peeled from the cavity with tweezers and suspended in 100 µl PBS with 0.5% FBS that was kept on ice. The suspended kidney was then homogenized by pipetting, vortexed briefly and then passed through a 35 µm cell strainer using a syringe plunger to help push the cells through into a microcentrifuge tube. The strainer was rinsed with extra buffer. Samples were kept on ice.

The viability stain FVS 510 (BD biosciences) was used for excluding dead cells from the analysis. Briefly, cells were suspended in 2 ml PBS, pelleted at 400 x g for 5 min. The supernatant was removed, and the pellet resuspended in 1 ml PBS. 1 µl of stain was added and the tubes were gently vortexed. Samples were incubated for 30 min. Cells were washed twice with 2 ml PBS with 0.5% FBS. Finally, cells were resuspended in PBS with 0.5% FBS. Just before sorting, the samples were passed through a strainer cap of a FACS tube by centrifugation to ensure that there were no clumps (400 x g, 00:30). The flow cytometry and sorting were done with FACS Arya Fusion (BD Biosciences). Cells were gated according to (95).

20,000 events were recorded for each sample. Sorted cells were kept on ice and pelleted after the flow cytometry was finished. RNA was extracted with a RNeasy Micro Plus Kit (Qiagen). Flow cytometry results were analysed with FlowJo 10.7.1.

To sort out neutrophils for RNA sequencing, fish homozygous for *pycard^tpu4^*and WT controls crossed to *Tg(mpx:GFP)i114* fish with fluorescent neutrophils were euthanized and dissected as described above. Dissected kidneys were suspended in 200 µl PBS with 1 % FBS and 25 µM HEPES and kept on ice. Suspended kidneys were homogenized by pipetting and vortexing briefly, before passing them through a 35 µm cell strainer with gentle (00:15-00:20) centrifugation. To exclude dead cells from the analysis, 5 µl of Propidium Iodide Staining Solution (Thermo Fisher Scientific) was added and samples were incubated for 5 minutes.

The flow cytometry and cell sorting were done with FACS Arya Fusion (BD Biosciences), based on fluorescence (GFP) emitted by neutrophils. Cells were gated according to Supplementary figure 10A. All events for each sample were recorded and sorted neutrophils were suspended to PBS with 1 % FBS and 25 µM HEPES. Cell suspensions were kept on ice and RNA was extracted with RNeasy Micro Plus Kit (Qiagen).

### Histology

8-month-old fish were injected with a low dose of *M. marinum* and euthanized 8wpi with an overdose of Tricaine. Fish were fixed in 10% phosphate-buffered formalin at room temperature in a rotator for 7 days after the removal of the heads and tails. Decalcification was carried out by incubating the samples for 7 days at room temperature in 0.5 M ethylenediaminetetraacetic acid (pH 8.0) in dH_2_O. Samples were incubated in 70% ethanol over night at room temperature with stirring after which they were carried through a rising ethanol series and transferred to xylene. Samples were then cast in paraffin and cut with a Leica microtome SM2010R starting from the dorsal side. Four 5 µm histological sections were collected at the intervals of 200 µm on StarFrost advanced adhesive (76 x 26 mm) glasses. Samples were deparaffinized and used for either Ziehl-Neelsen, trichrome or hypoxia staining. Ziehl-Neelsen staining was carried out according to a standard protocol and trichrome staining as described in Myllymäki et al. (2018) (28). After staining with Stainmate (Thermo Scientific), the slides were dehydrated with a series of ethanol solutions of increasing concentrations ending with xylene and embedded with DPX new (Sigma-Aldrich, Saint Louis, MO, USA). All slides were scanned with a Hamamatsu NanoZoomer S60 digital slide scanner (Hamamatsu, Hamamatsu City, Shizuoka Pref., Japan) and analysed with NDP View (Hamamatsu). Granulomas were counted and analysed based on their diameter, location, structure and hypoxicity.

Tissue hypoxia was detected with a Hypoxyprobe-1 kit (HP1-100Kit, Hypoxyprobe, Burlington, MA, USA). Pimonidazole hydrochloride (part of the HP1-100 kit) dissolved in PBS was injected intraperitoneally during terminal anaesthesia. Glasses were pre-treated with 0.05 M Tris – 0.01 M EDTA buffer with 0.05% Tween 20 (pH 9) using Lab Vision PT Module (Thermo Fisher Scientific). Endogenous peroxidase activity was blocked with a 5-minute incubation in 3% hydrogen peroxidase (VWR, 23614.291) and further by treating the samples with Bloxall blocking solution (SP-6000, Vector Laboratories, Burlingame, CA, USA) for 10 minutes. Samples were incubated with a 1:600 dilution of Hypoxyprobe-1 Mab1 (part of the HP1-100 kit) for 30 minutes after which they were treated with a secondary antibody, universal immune-peroxidase polymer anti-mouse complex (414131F, Nichirei Biosciences Inc., Tokyo, Japan), for 30 minutes. Staining was carried out with a 10-minute incubation with Histofine DAB-2V (425314F, Nichirei Biosciences) and counterstaining with a 2-minute incubation with Mayers Hematoxylin Plus (01825, Histolab Products AB, Askim, Sweden). An Autostainer 480 (Lab Vision, Thermo Fisher Scientific) was used to perform the staining. The glasses were dehydrated, embedded, and scanned as described above.

### mRNA sequencing

For RNA sequencing from a kidney (main hematopoietic organ in zebrafish), zebrafish were infected with a low dose of *M. marinum* or mock injected with PBS. At 4 wpi the fish were euthanised and the kidneys collected as explained for flow cytometry and cell sorting experiments. The kidneys were homogenized with ceramic beads as explained before, in the RNA preserving buffer RLT plus with β-mercaptoethanol. RNA was extracted with a RNeasy Plus Mini kit. RNA quality was assessed with a Fragment analyzer (Agilent, CA, USA). The RNA-sequencing service was bought from the Finnish Centre for Functional Genomics, Turku, Finland. The run was performed with NovaSeq 6000 S4 v1.5. (paired end sequencing, 100 bp read length, 20 million reads per sample depth). The RNA-seq data has been submitted to Gene expression omnibus (91,92) under the identifier GSE189627 (https://www.ncbi.nlm.nih.gov/geo/query/acc.cgi?acc=GSE189627).

The quality control of the RNA-sequencing read data was done using FastQC version 0.11.7 (96). The reads were aligned using STAR aligner (97) version 2.5.3a and the Ensembl reference genome GRCz11. Read counts for genes were quantified using featureCounts (98) version 1.6.2 and the Ensembl reference gene set release GRCz11.104 (99). A differential expression analysis was conducted using R version 3.6.1 (100) and DESeq2 (48) version 1.24.0. Differentially expressed genes were determined as genes that have a p-value <0.05 after adjustment for multiple testing, |log2 fold change| >1 and an absolute median difference of library-size normalized read counts >13 between two conditions. Gene expression heat maps were generated with pheatmap version 1.0.12. Differentially expressed genes were classified based on the data available from the Ensembl genome browser (101) version 104, Reactome (61), the Zebrafish Information Network ZFIN (102) and the literature.

For RNA sequencing from neutrophils, *pycard^tpu4/tpu4^*and WT controls (*pycard^tpu4^* originally crossed to transgenic *Tg(mpx:GFP)i114 (AB)* fish) were infected with *M. marinum* (mean dose 95, range 69-118 CFU). At 4 wpi, fish were euthanized, and kidney marrow-derived neutrophils were obtained as described for flow cytometry and cell sorting experiments. RNA from neutrophils was extracted with RNeasy Micro Plus Kit (Qiagen) and quality of RNA was assessed with NanoDrop (Thermo Fisher Scientific, Waltham, Massachusetts, USA).

The RNA sequencing service and bioinformatic analysis were bought from Novogene, Cambridge, United Kingdom. Library construction was performed on the Illumina platform (paired-end sequencing, 150 bp read length) yielding > 20 million reads per sample. The reads were aligned with using HISAT2 (103) and Ensembl reference genome GRz11, gene set release grcz11_gca_000002035_4 (99). Analysis of differentially expressed genes was performed with DESeq2 (49). The RNA sequencing data set has been submitted to Gene expression omnibus (91,92) under the identifier GSE270136 (https://www.ncbi.nlm.nih.gov/geo/query/acc.cgi?acc<u>= GSE270136</u>).

For further analysis, genes with *p* value < 0.05 before adjustment for multiple testing, at least two samples with ≥20 normalized reads, |log2 fold change| >1 and |fold change of medians| > 2 between the groups were included. The genes were studied using Ensembl BioMart (104,105) and were categorized according to Reactome (61). Reactome IDs were converted to Reactome pathway names using R version 4.3.2 (100) and reactome.db (106). Genes related to significantly up-and downregulated genes were classified based on the data available from the Ensembl genome browser (105) versions 111 and 112, gene ontology data obtained from BioMart (104,105), the Zebrafish Information Network ZFIN (102), GeneCards (107) and the literature.

### Imaging and quantifying the myeloid cells

The offspring of heterozygous *pycard^tpu4^* mutants carrying fluorescent neutrophils (homozygous mutants crossed to *Tg(mpx:GFP)*) or macrophages (homozygous mutants crossed to *Tg(mpeg:GFP)*) was dechorionated and transferred to embryonic medium supplemented with 0.0045% PTU (P7629, Sigma-Alrich) at 1 dpf. At 2 dpf the fluorescent larvae were picked using Nikon AZ100 macroscope (Nikon, Minato City, Tokyo, Japan). At 3 dpf the larvae were wounded under microscope with Fine-Ject 30G/0,3×12mm needle (4710003012, Henke Sass Wolf GmbH, Tuttlingen, Germany) on the side of the tail fin, embedded in 1.7% or 2% methyl cellulose (M0512-100G, Sigma-Aldrich) and imaged in one representative focus layer with Nikon AZ100 macroscope and NIS-Elements D 5.02.00 software (Nikon) 1.5 to 3 hours post wounding. The exposure time on the green channel was set to 600 ms or 1000 ms for 125 ms for the *Tg(mpx:GFP)* and Tg(*mpeg1.1:GFP)* line. The figures were saved as tiff. Fluorescent neutrophils and macrophages were manually calculated from unmodified figures with 120% or 150% magnification, respectively, using Corel PaintShop Pro 2020 version 22.0.0.132 (Alludo, Ottawa, Canada). The larvae were genotyped as described in the section “CRISPR-Cas9 mutagenesis and genotyping”.

### Power calculations and statistical analysis

Based on our previous data, we estimated the number of animals required for the survival and bacterial burden experiments (28,32,34,35,45). With the clincalc.com/stats/samplesize.aspx calculator, the required group size was estimated to be 16-20 fish (alpha 0.05, power 0.8). As the experiment was done blinded, a moderately higher group size of 25 was selected so that it contains a sufficient number of each genotype. Statistical analyses were done with Graph Pad prism 5.02. p-values below 0.05 were considered significant.

## Acknowledgments

We would like to thank Leena Mäkinen and Hannaleena Piippo for their assistance with the zebrafish experiments. In addition, we would like to thank Sari Toivola and Laura Kummola (Tampere University) for assistance in the histology and flow cytometry experiments, respectively, and Markus Ojanen for advice concerning flow cytometry experiments. We acknowledge the Tampere Histology Facility (HF), the Tampere Zebrafish Laboratory, Tampere facility of Protein Services (PS), Tampere University Flow Cytometry Facility (TFCF) and the Biocenter Finland (BF) and Tampere Genomics Facility for the service, Heini Huhtala for help with statistics and Helen Cooper for proof-reading the manuscript. We would like to thank the Finnish Functional Genomics Centre for their assistance in performing the RNA-seq analyses. We would like to thank the students who contributed to lab work: Riikka Penttinen, Mikko Kaasinen, Sarah Snoeck, Lien Kindt and Sini Saarimaa.

## Supplementary information captions

**Supplementary figure 1. Generation of *pycard* mutants with CRISPR-Cas9.**

A) The structure of the zebrafish *pycard* gene is presented with a box indicating the gRNA binding site. Exons 1-5 are indicated with e1-e5, introns (not drawn to scale) have been indicated with horizontal lines joining the exons, and the untranslated regions (3’UTR and 5’UTR) are indicated with a darker grey colour. The primers used in genotyping are indicated with a half arrow and a capital letter (F=forward primer, R=reverse primer), primers for transcript quantitation are indicated with lower case letter (f=forward primer, r=reverse primer). The reverse primer for quantitation is designed across the intron, as indicated by a dashed line over the intron. Below, the B) WT and C) the mutant *pycard^tpu4^*(left) and D) the mutant *pycard^tpu5^* DNA and coding sequences are shown underlined, with a box indicating the gRNA binding site and amino acids resulting from the frame shift are bolded. The length of the resulting amino acid chain is marked at the end. E-F) The gene expression was measured with qPCR in *pycard*^+/+^ and *pycard*^-/-^ larvae at 1, 3, 5 and 7 days postfertilization (dpf) for E) the mutant line *pycard^tpu4^* F) mutant line *pycard^tpu5^*. Results were normalized to *eef1α* expression. Each sample consists of 3-5 pooled larvae. The line indicates median.

**Supplementary figure 2. The *pycard^tpu5^* mutants are susceptible for *M. marinum* infection.**

Adult fish of the *pycard^tpu5^* line were infected with a low dose of *M. marinum* and their survival was followed daily. A) The first experiment, mean dose 14 CFU, range 9-18 CFU. End point survival proportions *pycard^+/+^*: 78.6%, *pycard^+/tpu5^*: 83.3%, *pycard^tpu5/tpu5^*: 50.0%. B) Mean dose 40 CFU, range 25-57 CFU. End point survival proportions *pycard^+/+^*: 86.4%, *pycard^+/tpu5^*: 82.6%, *pycard^tpu5/tpu5^*: 45.5%. The fish were genotyped post-mortem. The survival data are presented as a Kaplan-Meier survival curve. The statistical analysis was done with a log rank test. The second survival experiment b) was terminated earlier to minimize unnecessary suffering.

**Supplementary figure 3. *pycard*^-/-^ adult fish display a higher bacterial burden.**

Zebrafish from both mutant lines were infected with a low dose of *M. marinum*, and the bacterial burden was analysed at 4 wpi in whole organ block DNA using qPCR with *M. marinum* genome specific primers. A) *pycard^tpu4^*, medians for both groups *pycard^+/+^*: 34129 CFU, *pycar^tpu4/tpu4^*: 306007 CFU, p=0.049*. B) *pycard^tpu4^*, medians for both groups *pycard^+/+^*: 792 CFU, *pycard^tpu4/tpu4^*: 34612 CFU, p=0.006**. C) *pycard^tpu5^*, medians for both groups *pycard^+/+^*: 2636 CFU, *pycard^tpu5/tpu5^*: 10992 CFU, p=0.031*, D) *pycard^tpu5^*, *pycard^+/+^*: 167027 CFU, *pycard^tpu5/tpu5^*: 87447 CFU, p=0.340, E) *pycard^tpu5^*, *pycard^+/+^*: 120490 CFU, *pycard^tpu5/tpu5^*: 199455 CFU, p=0.187. The line indicates the median, the data were analysed with the Mann-Whitney U-test, two tailed. For statistical reasons, samples whose quantitation failed or was below 100 CFU detection limit (DL), a value of 100 CFU for *pycard*^+/+^ and 0 CFU for *pycard*^-/-^ were given.

**Supplementary figure 4. Flow cytometry gating strategy and results from individual experiments.**

A) Cells were gated from debris using forward scatter (FSC-A) vs. side scatter (SSC-A). Single cells were gated from all cells using FSC-A vs. FCS-H (pulse geometry gating). Live cells were gated based on the FVS510 live-dead stain. Leukocyte populations were gated based on size (FSC-A) and granularity (SSC-A). B) The experiment was repeated twice, and individual experiments are shown with either empty or filled symbols. 20 000 events were recorded per sample. Bacterial burden in experiment 1, mean dose 17 CFU, range 12.28 CFU. Bacterial burden in experiment 2, mean 19 CFU, range 16-21 CFU.

**Supplementary figure 5. Granuloma sizes per individual fish.**

Zebrafish of the mutant line *pycard^tpu5^* were infected with a low dose of *M. marinum* and at 8 wpi were sacrificed for analysis. All granulomas from each fish were analysed for their size. A linear mixed model was used for determining the effect of genotype on granuloma size (R-package lme4, fish as a random and genotype as a fixed factor). The lines indicate median and the interquartile range.

**Supplementary figure 6. Results from the granuloma characterization.**

Each granuloma from the fish from the *pycard^tpu5^* line was A) quantitated and characterised for whether it presented features of a B) nascent, C) necrotic, D) fibrous capsule or E) surrounding hypoxia. The fraction of each type from the total number of granulomas in a fish is presented for both genotypes. No significant differences were observed.

**Supplementary** figure 7**. Quantitative PCR analysis of selected genes in *pycard^tpu^*^5^ mutant line**.

Zebrafish were either mock injected or infected with a low dose of *M. marinum* (Mean dose 24 CFU, range 16-32 CFU). At 4 wpi, the fish were sacrificed and their kidney collected for RNA extraction. Transcripts were quantitated with qPCR. The line indicates the median.

**Supplementary figure 8. Analysis of the number of neutrophils and macrophages in *pycard^tpu4^* mutants at 3 dpf.**

*pycard^tpu4/tpu4^* zebrafish were crossed to transgenic *Tg(mpx:GFP)i114 (AB)* and *Tg(mpeg1.1:GFP)ka101 (AB)* zebrafish lines to receive *pycard^+/tpu4^* zebrafish with fluorescent macrophages or neutrophils, respectively. These fish were incrossed and at 3 dpi the progeny was wounded and and imaged with Nikon AZ100 macroscope and NIS-Elements D 5.02.00 software. The neutrophils and macrophages were manually counted from the figures using Corel PaintShop Pro 2020 version 22.0.0.132. The loss of *pycard* did not affect the number of A) neutrophils or B) macrophages in the larvae post wounding. The data in figure B) are pooled from two experiments. A two-tailed Mann-Whitney test was used for the statistical comparison of differences. C) Representative images are shown from each group. Brightness and contrast were adjusted to the same level for the larvae with fluorescent neutrophils and for the larvae with fluorescent macrophages, the channels were merged and the figures flipped and/or rotated if needed with Fiji (Image J) (90). The scale bar is 500 µm.

**Supplementary figure 9. Flow cytometry gating strategy, bacterial burden and 20 most expressed genes in WT fish derived neutrophils upon *M. marinum* infection (4 wpi).**

A) Cells were gated from debris using forward scatter (FSC-A) vs. side scatter (SSC-A). Single cells were gated from all cells using FSC-A vs. FCS-H (pulse geometry gating). Live cells were gated based on the Propidium Iodide Staining Solution. Neutrophil populations were gated based on fluorescence (GFP), size (FSC-A) and granularity (SSC-A). 20 000 events were recorded per sample. Bacterial dose in the experiment, mean 95 CFU, range 69-118 CFU. B) Percentage of neutrophils of total cell population. C) bacterial burden in *M. marinum* infected *pycard^+/+^* and *pycard^tpu4/tpu4^* fish (*pycard^tpu4^*line crossed to *Tg(mpx:GFP)i114* with fluorescent neutrophils). D) 20 most expressed genes in *pycard^+/+^* fish upon *M. marinum* infection (4 wpi) are dominantly expressed in neutrophils according to data by Tang et al. (2017) (51) and accessed via the online tool developed by Lareau et al. (2017) (52).

**Supplementary table 1: Differentially expressed genes in the adult *pycard^tpu4/tpu4^* zebrafish at basal state in PBS injected controls (4 wpi timepoint)**

**Supplementary table 2: Upregulated genes in the adult *pycard^tpu4/tpu4^* zebrafish infected with *M. marinum* (4 wpi timepoint)**

**Supplementary table 3: Downregulated genes in the adult *pycard^tpu4/tpu4^* zebrafish infected with *M. marinum* (4 wpi timepoint)**

**Supplementary table 4: Differentially expressed genes in neutrophils of adult *pycard^tpu4/tpu4^* zebrafish infected with *M. marinum* (4 wpi timepoint) , P < 0.05**

**Supplementary table 5: Differentially expressed genes in neutrophils of adult *pycard^tpu4/tpu4^* zebrafish infected with *M. marinum* (4 wpi timepoint) categorized according to Reactome**

**Supplementary table 6: Upregulated genes in neutrophils of adult *pycard^tpu4/tpu4^* zebrafish infected with *M. marinum* (4 wpi timepoint)**

**Supplementary table 7: Downregulated genes in neutrophils of adult *pycard^tpu4/tpu4^* zebrafish infected with *M. marinum* (4 wpi timepoint)**

## References

1. Global Tuberculosis Report 2023 [Internet]. [cited 2024 Jun 7]. Available from: https://www.who.int/teams/global-tuberculosis-programme/tb-reports/global-tuberculosis-report-2023

2. Andersen P, Doherty TM. The success and failure of BCG - implications for a novel tuberculosis vaccine. Nature Reviews Microbiology. 2005;3(8):656–62.

3. Furin J, Cox H, Pai M. Tuberculosis. Lancet. 2019 Apr 20;393(10181):1642–56.

4. Houben RMGJ, Dodd PJ. The Global Burden of Latent Tuberculosis Infection: A Re-estimation Using Mathematical Modelling. Metcalfe JZ, editor. PLoS Med. 2016 Oct 25;13(10):e1002152.

5. Ai JW, Ruan QL, Liu QH, Zhang WH. Updates on the risk factors for latent tuberculosis reactivation and their managements. Emerging Microbes & Infections. 2016 Jan 1;5(1):1–8.

6. Churchyard G, Kim P, Shah NS, Rustomjee R, Gandhi N, Mathema B, et al. What We Know About Tuberculosis Transmission: An Overview. The Journal of Infectious Diseases. 2017 Nov 3;216(suppl_6):S629–35.

7. Goren MB, Hart PD, Young MR, Armstrong JA. Prevention of phagosome-lysosome fusion in cultured macrophages by sulfatides of Mycobacterium tuberculosis. PNAS. 1976 Jul 1;73(7):2510–4.

8. Goldberg MF, Saini NK, Porcelli SA. Evasion of Innate and Adaptive Immunity by Mycobacterium tuberculosis. Microbiol Spectr. 2014 Oct;2(5).

9. Korb VC, Chuturgoon AA, Moodley D. Mycobacterium tuberculosis: Manipulator of Protective Immunity. International Journal of Molecular Sciences. 2016 Mar;17(3):131.

10. Master SS, Rampini SK, Davis AS, Keller C, Ehlers S, Springer B, et al. Mycobacterium tuberculosis Prevents Inflammasome Activation. Cell host & microbe. 2008;3(4):224–32.

11. Rastogi S, Ellinwood S, Augenstreich J, Mayer-Barber KD, Briken V. Mycobacterium tuberculosis inhibits the NLRP3 inflammasome activation via its phosphokinase PknF. PLoS Pathog. 2021 Jul;17(7):e1009712.

12. Martinon F, Burns K, Rg Tschopp J. The Inflammasome: A Molecular Platform Triggering Activation of Inflammatory Caspases and Processing of proIL-␤ that they possess several distinct protein/protein inter-action domains which are used to assemble large multi-component complexes. Apaf-1, for example, contains an N-terminal CARD followed by a NBS/self-oligomerization domain and a C-terminal WD-40 repeat (Jaro. Molecular Cell. 2002;10(Generic):417.

13. Bergsbaken T, Fink SL, Cookson BT. Pyroptosis: host cell death and inflammation. Nat Rev Microbiol. 2009;7(2):99–109.

14. Mayer-Barber KD, Barber DL, Shenderov K, White SD, Wilson MS, Cheever A, et al. Caspase-1 independent IL-1beta production is critical for host resistance to mycobacterium tuberculosis and does not require TLR signaling in vivo. J Immunol. 2010 Apr 1;184(7):3326–30.

15. McElvania Tekippe E, Allen IC, Hulseberg PD, Sullivan JT, McCann JR, Sandor M, et al. Granuloma formation and host defense in chronic Mycobacterium tuberculosis infection requires PYCARD/ASC but not NLRP3 or caspase-1. PLoS One. 2010 Aug 20;5(8):e12320.

16. Mishra BB, Moura-Alves P, Sonawane A, Hacohen N, Griffiths G, Moita LF, et al. Mycobacterium tuberculosis protein ESAT-6 is a potent activator of the NLRP3/ASC inflammasome. Cell Microbiol. 2010 Aug;12(8):1046–63.

17. Beckwith KS, Beckwith MS, Ullmann S, Sætra RS, Kim H, Marstad A, et al. Plasma membrane damage causes NLRP3 activation and pyroptosis during Mycobacterium tuberculosis infection. Nat Commun. 2020 May 8;11(1):2270.

18. Dorhoi A, Nouailles G, Jörg S, Hagens K, Heinemann E, Pradl L, et al. Activation of the NLRP3 inflammasome by Mycobacterium tuberculosis is uncoupled from susceptibility to active tuberculosis. European Journal of Immunology. 2012;42(2):374–84.

19. Javanmard Khameneh H, Leong KWK, Mencarelli A, Vacca M, Mambwe B, Neo K, et al. The Inflammasome Adaptor ASC Intrinsically Limits CD4+ T-Cell Proliferation to Help Maintain Intestinal Homeostasis. Front Immunol [Internet]. 2019;10(Journal Article). Available from: https://www.frontiersin.org/articles/10.3389/fimmu.2019.01566/full

20. Taxman DJ, Holley-Guthrie EA, Huang MTH, Moore CB, Bergstralh DT, Allen IC, et al. The NLR adaptor ASC/PYCARD regulates DUSP10, mitogen-activated protein kinase (MAPK), and chemokine induction independent of the inflammasome. J Biol Chem. 2011 Jun 3;286(22):19605–16.

21. Taxman DJ, Zhang J, Champagne C, Bergstralh DT, Iocca HA, Lich JD, et al. Cutting edge: ASC mediates the induction of multiple cytokines by Porphyromonas gingivalis via caspase-1-dependent and -independent pathways. J Immunol. 2006 Oct 1;177(7):4252–6.

22. Ippagunta SK, Malireddi RKS, Shaw PJ, Neale GA, Vande Walle L, Green DR, et al. The inflammasome adaptor ASC regulates the function of adaptive immune cells by controlling Dock2-mediated Rac activation and actin polymerization. Nat Immunol. 2011 Sep 4;12(10):1010–6.

23. Uchiyama R, Yonehara S, Taniguchi S, Ishido S, Ishii KJ, Tsutsui H. Inflammasome and Fas-Mediated IL-1β Contributes to Th17/Th1 Cell Induction in Pathogenic Bacterial Infection In Vivo. JI. 2017;199(3):1122.

24. Lohi O, Parikka M, Rämet M. The zebrafish as a model for paediatric diseases. Acta Paediatrica. 2013 Feb;102(2):104–10.

25. Howe K, Torroja CF, Torrance J, Collins JE, Humphray S, McLaren K, et al. The zebrafish reference genome sequence and its relationship to the human genome. Nature (London). 2013;496(7446):498–503.

26. Cronan MR, Tobin DM. Fit for consumption: zebrafish as a model for tuberculosis. Disease models & mechanisms. 2014;7(7):777–84.

27. Meijer AH. Protection and pathology in TB: learning from the zebrafish model. Semin Immunopathol. 2016 Mar 1;38(2):261–73.

28. Myllymäki H, Niskanen M, Luukinen H, Parikka M, Rämet M. Identification of protective postexposure mycobacterial vaccine antigens using an immunosuppression-based reactivation model in the zebrafish. Dis Model Mech. 2018 Mar 13;11(3):dmm033175.

29. Myllymaki H, Bauerlein CA, Ramet M. The Zebrafish Breathes new Life into the Study of Tuberculosis. Frontiers in Immunology. 2016;7(Journal Article):196.

30. Tobin DM, May RC, Wheeler RT. Zebrafish: A See-Through Host and a Fluorescent Toolbox to Probe Host–Pathogen Interaction. PLOS Pathogens. 2012 Jan 5;8(1):e1002349.

31. Tobin DM, Ramakrishnan L. Comparative pathogenesis of Mycobacterium marinum and Mycobacterium tuberculosis. CellMicrobiol. 2008;10(5):1027–39.

32. Parikka M, Hammaren MM, Harjula SKE, Halfpenny NJA, Oksanen KE, Lahtinen MJ, et al. Mycobacterium marinum Causes a Latent Infection that Can Be Reactivated by Gamma Irradiation in Adult Zebrafish. Plos Pathogens. 2012;8(9).

33. Swaim LE, Connolly LE, Volkman HE, Humbert O, Born DE, Ramakrishnan L. Mycobacterium marinum infection of adult zebrafish causes caseating granulomatous tuberculosis and is moderated by adaptive immunity (vol 74, pg 6108, 2006). InfectImmun. 2007;75(3):1540.

34. Ojanen MJT, Turpeinen H, Cordova ZM, Hammarén MM, Harjula SKE, Parikka M, et al. The Proprotein Convertase Subtilisin/Kexin FurinA Regulates Zebrafish Host Response against Mycobacterium marinum. Ehrt S, editor. Infect Immun. 2015 Apr;83(4):1431–42.

35. Harjula SKE, Ojanen MJT, Taavitsainen S, Nykter M, Rämet M. Interleukin 10 mutant zebrafish have an enhanced interferon gamma response and improved survival against a Mycobacterium marinum infection. Sci Rep. 2018 Jul 9;8(1):10360.

36. Ojanen MJT, Uusi-Mäkelä MIE, Harjula SKE, Saralahti AK, Oksanen KE, Kähkönen N, et al. Intelectin 3 is dispensable for resistance against a mycobacterial infection in zebrafish (Danio rerio). Sci Rep. 2019 Jan 30;9(1):995.

37. Lam SH, Chua HL, Gong Z, Lam TJ, Sin YM. Development and maturation of the immune system in zebrafish, Danio rerio: a gene expression profiling, in situ hybridization and immunological study. Developmental & Comparative Immunology. 2004 Jan 1;28(1):9–28.

38. Page DM, Wittamer V, Bertrand JY, Lewis KL, Pratt DN, Delgado N, et al. An evolutionarily conserved program of B-cell development and activation in zebrafish. Blood. 2013 Aug 22;122(8):e1–11.

39. Frame JM, Kubaczka C, Long TL, Esain V, Soto RA, Hachimi M, et al. Metabolic Regulation of Inflammasome Activity Controls Embryonic Hematopoietic Stem and Progenitor Cell Production. Developmental Cell. 2020 Oct 26;55(2):133–149.e6.

40. Kuri P, Schieber NL, Thumberger T, Wittbrodt J, Schwab Y, Leptin M. Dynamics of in vivo ASC speck formation. J Cell Biol. 2017 Sep 4;216(9):2891–909.

41. Li JY, Wang YY, Shao T, Fan DD, Lin AF, Xiang LX, et al. The zebrafish NLRP3 inflammasome has functional roles in ASC-dependent interleukin-1β maturation and gasdermin E–mediated pyroptosis. J Biol Chem. 2019;295(4):1120.

42. Tyrkalska SD, Candel S, Angosto D, Gómez-Abellán V, Martín-Sánchez F, García-Moreno D, et al. Neutrophils mediate Salmonella Typhimurium clearance through the GBP4 inflammasome-dependent production of prostaglandins. Nat Commun. 2016 Jul 1;7(1):12077.

43. Tyrkalska SD, Candel S, Pérez-Oliva AB, Valera A, Alcaraz-Pérez F, García-Moreno D, et al. Identification of an Evolutionarily Conserved Ankyrin Domain-Containing Protein, Caiap, Which Regulates Inflammasome-Dependent Resistance to Bacterial Infection. Front Immunol. 2017;8:1375.

44. Tyrkalska SD, Pérez-Oliva AB, Rodríguez-Ruiz L, Martínez-Morcillo FJ, Alcaraz-Pérez F, Martínez-Navarro FJ, et al. Inflammasome Regulates Hematopoiesis through Cleavage of the Master Erythroid Transcription Factor GATA1. Immunity. 2019 Jul 16;51(1):50–63.e5.

45. Harjula SKE, Saralahti AK, Ojanen MJT, Rantapero T, Uusi-Mäkelä MIE, Nykter M, et al. Characterization of immune response against Mycobacterium marinum infection in the main hematopoietic organ of adult zebrafish (Danio rerio). Developmental and comparative immunology [Internet]. 2020;103(Journal Article). Available from: 10.1016/j.dci.2019.103523

46. Matty MA, Knudsen DR, Walton EM, Beerman RW, Cronan MR, Pyle CJ, et al. Potentiation of P2RX7 as a host-directed strategy for control of mycobacterial infection. Elife. 2019 Jan 29;8:e39123.

47. Yan S, Shen H, Lian Q, Jin W, Zhang R, Lin X, et al. Deficiency of the AIM2-ASC Signal Uncovers the STING-Driven Overreactive Response of Type I IFN and Reciprocal Depression of Protective IFN-γ Immunity in Mycobacterial Infection. J Immunol. 2018 Feb 1;200(3):1016–26.

48. Love MI, Huber W, Anders S. Moderated estimation of fold change and dispersion for RNA-seq data with DESeq2. Genome Biol. 2014;15(12):550.

49. Anders S, Huber W. Differential expression analysis for sequence count data. Genome Biol. 2010 Oct;11(10):R106.

50. Carmona SJ, Teichmann SA, Ferreira L, Macaulay IC, Stubbington MJT, Cvejic A, et al. Single-cell transcriptome analysis of fish immune cells provides insight into the evolution of vertebrate immune cell types. Genome Res. 2017 Mar;27(3):451–61.

51. Tang Q, Iyer S, Lobbardi R, Moore JC, Chen H, Lareau C, et al. Dissecting hematopoietic and renal cell heterogeneity in adult zebrafish at single-cell resolution using RNA sequencing. J Exp Med. 2017 Oct 2;214(10):2875–87.

52. Lareau C, Iyer S, Langenau DM, Aryee M. Single Cell inDrops RNA-Seq Visualization of Adult Zebrafish Whole Kidney Marrow [Internet]. Harvard University; 2017. Available from: https://molpath.shinyapps.io/zebrafishblood/

53. Gieger C, Radhakrishnan A, Cvejic A, Tang W, Porcu E, Pistis G, et al. New gene functions in megakaryopoiesis and platelet formation. Nature. 2011 Dec;480(7376):201–8.

54. Zhang Y, Xue Y, Cao C, Huang J, Hong Q, Hai T, et al. Thyroid hormone regulates hematopoiesis via the TR-KLF9 axis. Blood. 2017 Nov 16;130(20):2161–70.

55. Jung J, Buisman SC, Weersing E, Dethmers-Ausema A, Zwart E, Schepers H, et al. CBX7 Induces Self-Renewal of Human Normal and Malignant Hematopoietic Stem and Progenitor Cells by Canonical and Non-canonical Interactions. Cell Reports. 2019 Feb 12;26(7):1906–1918.e8.

56. Zhu J, Zhang Y, Joe GJ, Pompetti R, Emerson SG. NF-Ya activates multiple hematopoietic stem cell (HSC) regulatory genes and promotes HSC self-renewal. Proc Natl Acad Sci U S A. 2005 Aug 16;102(33):11728–33.

57. Watamoto K, Towatari M, Ozawa Y, Miyata Y, Okamoto M, Abe A, et al. Altered interaction of HDAC5 with GATA-1 during MEL cell differentiation. Oncogene. 2003 Dec 11;22(57):9176–84.

58. Kalev-Zylinska ML, Horsfield JA, Flores MVC, Postlethwait JH, Chau JYM, Cattin PM, et al. Runx3 is required for hematopoietic development in zebrafish. Dev Dyn. 2003 Nov;228(3):323–36.

59. Espina J, Feijóo CG, Solís C, Glavic A. csrnp1a Is Necessary for the Development of Primitive Hematopoiesis Progenitors in Zebrafish. PLOS ONE. 2013 Jan 9;8(1):e53858.

60. Segovia M, Russo S, Jeldres M, Mahmoud YD, Perez V, Duhalde M, et al. Targeting TMEM176B Enhances Antitumor Immunity and Augments the Efficacy of Immune Checkpoint Blockers by Unleashing Inflammasome Activation. Cancer Cell. 2019 May 13;35(5):767–781.e6.

61. Griss J, Viteri G, Sidiropoulos K, Nguyen V, Fabregat A, Hermjakob H. ReactomeGSA - Efficient Multi-Omics Comparative Pathway Analysis. Molecular & Cellular Proteomics. 2020 Dec 1;19(12):2115–25.

62. Zhao W, Shi CS, Harrison K, Hwang IY, Nabar NR, Wang M, et al. AKT Regulates NLRP3 Inflammasome Activation by Phosphorylating NLRP3 Serine 5. The Journal of Immunology. 2020 Oct 15;205(8):2255–64.

63. Zhang BH, Liu H, Yuan Y, Weng XD, Du Y, Chen H, et al. Knockdown of TRIM8 Protects HK-2 Cells Against Hypoxia/Reoxygenation-Induced Injury by Inhibiting Oxidative Stress-Mediated Apoptosis and Pyroptosis via PI3K/Akt Signal Pathway. DDDT. 2021 Dec;Volume 15:4973–83.

64. Zhu J, Dong J, Ji L, Jiang P, Leung TF, Liu D, et al. Anti-Allergic Inflammatory Activity of Interleukin-37 Is Mediated by Novel Signaling Cascades in Human Eosinophils. Front Immunol. 2018 Jun 22;9:1445.

65. Hu Z, Xuan L, Wu T, Jiang N, Liu X, Chang J, et al. Taxifolin attenuates neuroinflammation and microglial pyroptosis via the PI3K/Akt signaling pathway after spinal cord injury. International Immunopharmacology. 2023 Jan;114:109616.

66. Li Y, Huang Y, Cao X, Yin X, Jin X, Liu S, et al. Functional and structural characterization of zebrafish ASC. FEBS J. 2018 Jul;285(14):2691–707.

67. Cheong M, Gartlan KH, Lee JS, Tey SK, Zhang P, Kuns RD, et al. ASC Modulates CTL Cytotoxicity and Transplant Outcome Independent of the Inflammasome. Cancer Immunol Res. 2020 Aug;8(8):1085–98.

68. Pedrosa J, Saunders BM, Appelberg R, Orme IM, Silva MT, Cooper AM. Neutrophils Play a Protective Nonphagocytic Role in Systemic Mycobacterium tuberculosis Infection of Mice. Infection and Immunity. 2000 Feb 1;68(2):577–83.

69. Sugawara I, Udagawa T, Yamada H. Rat Neutrophils Prevent the Development of Tuberculosis. Infect Immun. 2004 Mar;72(3):1804–6.

70. Kroon EE, Coussens AK, Kinnear C, Orlova M, Möller M, Seeger A, et al. Neutrophils: Innate Effectors of TB Resistance? Front Immunol. 2018 Nov 14;9:2637.

71. Kenyon A, Gavriouchkina D, Zorman J, Napolitani G, Cerundolo V, Sauka-Spengler T. Active nuclear transcriptome analysis reveals inflammasome-dependent mechanism for early neutrophil response to Mycobacterium marinum. Sci Rep. 2017 Jul 26;7(1):6505.

72. Zhao JL, Ma C, O’Connell RM, Mehta A, DiLoreto R, Heath JR, et al. Conversion of Danger Signals into Cytokine Signals by Hematopoietic Stem and Progenitor Cells for Regulation of Stress-Induced Hematopoiesis. Cell Stem Cell. 2014 Apr 3;14(4):445–59.

73. Hawkins PT, Stephens LR, Suire S, Wilson M. PI3K Signaling in Neutrophils. In: Rommel C, Vanhaesebroeck B, Vogt PK, editors. Phosphoinositide 3-kinase in Health and Disease [Internet]. Berlin, Heidelberg: Springer Berlin Heidelberg; 2010 [cited 2024 May 2]. p. 183–202. (Current Topics in Microbiology and Immunology; vol. 346). Available from: http://link.springer.com/10.1007/82_2010_40

74. Manning BD, Toker A. AKT/PKB Signaling: Navigating the Network. Cell. 2017 Apr;169(3):381– 405.

75. Choorapoikayil S, Kers R, Herbomel P, Kissa K, Den Hertog J. Pivotal role of Pten in the balance between proliferation and differentiation of hematopoietic stem cells in zebrafish. Blood. 2014 Jan 9;123(2):184–90.

76. Lozano-Gil JM, Rodríguez-Ruiz L, Tyrkalska SD, García-Moreno D, Pérez-Oliva AB, Mulero V. Gasdermin E mediates pyroptotic cell death of neutrophils and macrophages in a zebrafish model of chronic skin inflammation. Developmental & Comparative Immunology. 2022 Jul;132:104404.

77. Cirillo SLG, Subbian S, Chen B, Weisbrod TR, Jacobs WR, Cirillo JD. Protection of Mycobacterium tuberculosis from Reactive Oxygen Species Conferred by the mel2 Locus Impacts Persistence and Dissemination. Infection and Immunity. 2009 Jun 1;77(6):2557–67.

78. Kim SR, Lee YC, Kim HJ, Kim SH. Activation of NLRP3 inflammasome is regulated by mitochondrial ROS via PI3K-HIF-VEGF pathway in acute lung injury. European Respiratory Journal [Internet]. 2015 Sep 1 [cited 2021 Nov 24];46(suppl 59). Available from: https://erj.ersjournals.com/content/46/suppl_59/PA3026

79. Lu YX, Ju HQ, Liu ZX, Chen DL, Wang Y, Zhao Q, et al. ME1 Regulates NADPH Homeostasis to Promote Gastric Cancer Growth and Metastasis. Cancer Res. 2018 Apr 15;78(8):1972–85.

80. Noubade R, Wong K, Ota N, Rutz S, Eidenschenk C, Valdez PA, et al. NRROS negatively regulates reactive oxygen species during host defence and autoimmunity. Nature. 2014 May;509(7499):235–9.

81. Wu S, Xue R, Hassan S, Nguyen TML, Wang T, Pan H, et al. Il34-Csf1r Pathway Regulates the Migration and Colonization of Microglial Precursors. Dev Cell. 2018 Sep 10;46(5):552–563.e4.

82. Popovic M, Yaparla A, Paquin-Proulx D, Koubourli DV, Webb R, Firmani M, et al. Colony-stimulating factor-1- and interleukin-34-derived macrophages differ in their susceptibility to Mycobacterium marinum. Journal of Leukocyte Biology. 2019;106(6):1257–69.

83. Tobin DM, Vary JC, Ray JP, Walsh GS, Dunstan SJ, Bang ND, et al. The lta4h locus modulates susceptibility to mycobacterial infection in zebrafish and humans. Cell. 2010 Mar 5;140(5):717–30.

84. Tobin DM, Roca FJ, Ray JP, Ko DC, Ramakrishnan L. An Enzyme That Inactivates the Inflammatory Mediator Leukotriene B4 Restricts Mycobacterial Infection. PLOS ONE. 2013 Jul 11;8(7):e67828.

85. Stenger S, Rosat JP, Bloom BR, Krensky AM, Modlin RL. Granulysin: a lethal weapon of cytolytic T cells. Immunology Today. 1999 Sep 1;20(9):390–4.

86. Hortle E, Johnson KE, Johansen MD, Nguyen T, Shavit JA, Britton WJ, et al. Thrombocyte Inhibition Restores Protective Immunity to Mycobacterial Infection in Zebrafish. The Journal of Infectious Diseases. 2019 Jul 2;220(3):524–34.

87. Fox KA, Kirwan DE, Whittington AM, Krishnan N, Robertson BD, Gilman RH, et al. Platelets Regulate Pulmonary Inflammation and Tissue Destruction in Tuberculosis. Am J Respir Crit Care Med. 2018 Jul 15;198(2):245–55.

88. Kirwan DE, Chong DLW, Friedland JS. Platelet Activation and the Immune Response to Tuberculosis. Frontiers in Immunology. 2021;12:1871.

89. Eichelberger KR, Goldman WE. Manipulating neutrophil degranulation as a bacterial virulence strategy. Coers J, editor. PLoS Pathog. 2020 Dec 10;16(12):e1009054.

90. Geng X, Wu X, Yang Q, Xin H, Zhang B, Wang D, et al. Whole transcriptome sequencing reveals neutrophils’ transcriptional landscape associated with active tuberculosis. Front Immunol. 2022 Aug 18;13:954221.

91. Barrett T, Wilhite SE, Ledoux P, Evangelista C, Kim IF, Tomashevsky M, et al. NCBI GEO: archive for functional genomics data sets—update. Nucleic Acids Research. 2013 Jan 1;41(D1):D991–5.

92. Edgar R, Domrachev M, Lash AE. Gene Expression Omnibus: NCBI gene expression and hybridization array data repository. Nucleic Acids Res. 2002 Jan 1;30(1):207–10.

93. Renshaw SA, Loynes CA, Trushell DMI, Elworthy S, Ingham PW, Whyte MKB. A transgenic zebrafish model of neutrophilic inflammation. Blood. 2006 Dec 15;108(13):3976–8.

94. Tang R, Dodd A, Lai D, McNabb WC, Love DR. Validation of zebrafish (Danio rerio) reference genes for quantitative real-time RT-PCR normalization. Acta Biochim Biophys Sin (Shanghai). 2007;39(5):384–90.

95. Langenau DM, Ferrando AA, Traver D, Kutok JL, Hezel JPD, Kanki JP, et al. In vivo tracking of T cell development, ablation, and engraftment in transgenic zebrafish. PNAS. 2004 May 11;101(19):7369–74.

96. Andrews S. Fastqc. a quality control tool for high throughput sequence data. Vol. 2017. 2010.

97. Dobin A, Davis CA, Schlesinger F, Drenkow J, Zaleski C, Jha S, et al. STAR: ultrafast universal RNA-seq aligner. Bioinformatics. 2013 Jan 1;29(1):15–21.

98. Liao Y, Smyth GK, Shi W. featureCounts: an efficient general purpose program for assigning sequence reads to genomic features. Bioinformatics. 2014 Apr 1;30(7):923–30.

99. Hubbard T, Barker D, Birney E, Cameron G, Chen Y, Clark L, et al. The Ensembl genome database project. Nucleic Acids Research. 2002 Jan 1;30(1):38–41.

100. R: The R Project for Statistical Computing [Internet]. [cited 2024 May 15]. Available from: https://www.r-project.org/

101. Howe KL, Achuthan P, Allen J, Allen J, Alvarez-Jarreta J, Amode MR, et al. Ensembl 2021. Nucleic Acids Research. 2021 Jan 8;49(D1):D884–91.

102. Ruzicka L, Howe DG, Ramachandran S, Toro S, Van Slyke CE, Bradford YM, et al. The Zebrafish Information Network: new support for non-coding genes, richer Gene Ontology annotations and the Alliance of Genome Resources. Nucleic Acids Research. 2019 Jan 8;47(D1):D867–73.

103. Mortazavi A, Williams BA, McCue K, Schaeffer L, Wold B. Mapping and quantifying mammalian transcriptomes by RNA-Seq. Nat Methods. 2008 Jul;5(7):621–8.

104. Kinsella RJ, Kahari A, Haider S, Zamora J, Proctor G, Spudich G, et al. Ensembl BioMarts: a hub for data retrieval across taxonomic space. Database. 2011 Jul 23;2011(0):bar030–bar030.

105. Martin FJ, Amode MR, Aneja A, Austine-Orimoloye O, Azov AG, Barnes I, et al. Ensembl 2023. Nucleic Acids Research. 2023 Jan 6;51(D1):D933–41.

106. Ligtenberg W. reactome.db [Internet]. [object Object]; 2017 [cited 2024 May 15]. Available from: https://bioconductor.org/packages/reactome.db

107. Stelzer G, Rosen N, Plaschkes I, Zimmerman S, Twik M, Fishilevich S, et al. The GeneCards Suite: From Gene Data Mining to Disease Genome Sequence Analyses. CP in Bioinformatics [Internet]. 2016 Jun [cited 2024 May 15];54(1). Available from: https://currentprotocols.onlinelibrary.wiley.com/doi/10.1002/cpbi.5

108. Chestnut B, Sumanas S. Zebrafish etv2 knock-in line labels vascular endothelial and blood progenitor cells. Dev Dyn. 2020 Feb;249(2):245–61.

109. Xin GY, Li WG, Suman TY, Jia PP, Ma YB, Pei DS. Gut bacteria Vibrio sp. and Aeromonas sp. trigger the expression levels of proinflammatory cytokine: First evidence from the germ-free zebrafish. Fish & Shellfish Immunology. 2020 Nov 1;106:518–25.

110. Boudinot P, van der Aa LM, Jouneau L, Du Pasquier L, Pontarotti P, Briolat V, et al. Origin and evolution of TRIM proteins: new insights from the complete TRIM repertoire of zebrafish and pufferfish. PLoS One. 2011;6(7):e22022.

111. Climer LK, Cox AM, Reynolds TJ, Simmons DD. Oncomodulin: The Enigmatic Parvalbumin Protein. Front Mol Neurosci. 2019;12:235.

112. Liu X, Cao X, Wang S, Ji G, Zhang S, Li H. Identification of Ly2 members as antimicrobial peptides from zebrafish Danio rerio. Biosci Rep. 2017 Feb 28;37(1):BSR20160265.

113. Kobayashi I, Kondo M, Yamamori S, Kobayashi-Sun J, Taniguchi M, Kanemaru K, et al. Enrichment of hematopoietic stem/progenitor cells in the zebrafish kidney. Sci Rep. 2019 Oct 2;9(1):14205.

114. Somoza R, Beutler E. Phosphoglycolate phosphatase and 2,3-diphosphoglycerate in red cells of normal and anemic subjects. Blood. 1983 Oct;62(4):750–3.

115. Tokuhisa M, Kadowaki T, Ogawa K, Yamaguchi Y, Kido MA, Gao W, et al. Expression and localisation of Rab44 in immune-related cells change during cell differentiation and stimulation. Sci Rep. 2020 Jul 1;10(1):10728.

116. Nishio H, Suda T, Sawada K, Miyamoto T, Koike T, Yamaguchi Y. Molecular cloning of cDNA encoding human Rab3D whose expression is upregulated with myeloid differentiation. Biochim Biophys Acta. 1999 Feb 16;1444(2):283–90.

117. Foulkes MJ, Henry KM, Rougeot J, Hooper-Greenhill E, Loynes CA, Jeffrey P, et al. Expression and regulation of drug transporters in vertebrate neutrophils. Sci Rep. 2017 Jul 10;7(1):4967.

118. Pires D, Marques J, Pombo JP, Carmo N, Bettencourt P, Neyrolles O, et al. Role of Cathepsins in Mycobacterium tuberculosis Survival in Human Macrophages. Sci Rep. 2016 Aug 30;6(1):32247.

119. Jing W, Gershan JA, Holzhauer S, Weber J, Palen K, McOlash L, et al. T Cells Deficient in Diacylglycerol Kinase ζ Are Resistant to PD-1 Inhibition and Help Create Persistent Host Immunity to Leukemia. Cancer Res [Internet]. 2017 Sep 15 [cited 2021 Nov 30]; Available from: https://cancerres.aacrjournals.org/content/early/2017/10/02/0008-5472.CAN-17-1309

120. Kortum AN, Rodriguez-Nunez I, Yang J, Shim J, Runft D, O’Driscoll ML, et al. Differential expression and ligand binding indicate alternative functions for zebrafish polymeric immunoglobulin receptor (pIgR) and a family of pIgR-like (PIGRL) proteins. Immunogenetics. 2014 Apr;66(4):267–79.

121. Veneman WJ, Stockhammer OW, de Boer L, Zaat SAJ, Meijer AH, Spaink HP. A zebrafish high throughput screening system used for Staphylococcus epidermidis infection marker discovery. BMC Genomics. 2013 Apr 15;14:255.

122. Denisenko E, Guler R, Mhlanga M, Suzuki H, Brombacher F, Schmeier S. Transcriptionally induced enhancers in the macrophage immune response to Mycobacterium tuberculosis infection. BMC Genomics. 2019 Jan 22;20:71.

123. Akl I, Lelubre C, Uzureau P, Piagnerelli M, Biston P, Rousseau A, et al. Apolipoprotein L Expression Correlates with Neutrophil Cell Death in Critically Ill Patients. Shock. 2017 Jan;47(1):111–8.

124. Zhou Z jun, Sun L. Edwardsiella tarda-Induced Inhibition of Apoptosis: A Strategy for Intracellular Survival. Frontiers in Cellular and Infection Microbiology. 2016;6:76.

125. Chatzopoulou A, Heijmans JPM, Burgerhout E, Oskam N, Spaink HP, Meijer AH, et al. Glucocorticoid-Induced Attenuation of the Inflammatory Response in Zebrafish. Endocrinology. 2016 Jul;157(7):2772–84.

126. Cao M, Shikama Y, Kimura H, Noji H, Ikeda K, Ono T, et al. Mechanisms of Impaired Neutrophil Migration by MicroRNAs in Myelodysplastic Syndromes. The Journal of Immunology. 2017 Mar 1;198(5):1887–99.

127. Xu S, Xie F, Tian L, Manno SH, Manno FAM, Cheng SH. Prolonged neutrophil retention in the wound impairs zebrafish heart regeneration after cryoinjury. Fish Shellfish Immunol. 2019 Nov;94:447–54.

128. Bekpen C, Hunn JP, Rohde C, Parvanova I, Guethlein L, Dunn DM, et al. The interferon-inducible p47 (IRG) GTPases in vertebrates: loss of the cell autonomous resistance mechanism in the human lineage. Genome Biol. 2005;6(11):R92.

129. Rong C, Shi Y, Huang J, Wang X, Shimizu R, Mori Y, et al. The Effect of Metadherin on NF-κB Activation and Downstream Genes in Ovarian Cancer. Cell Transplant. 2020 Dec;29:963689720905506.

130. Chambers KF, Day PE, Aboufarrag HT, Kroon PA. Polyphenol Effects on Cholesterol Metabolism via Bile Acid Biosynthesis, CYP7A1: A Review. Nutrients. 2019 Oct 28;11(11):E2588.

131. Lu X, Nannenga B, Donehower LA. PPM1D dephosphorylates Chk1 and p53 and abrogates cell cycle checkpoints. Genes Dev. 2005 May 15;19(10):1162–74.

132. Bayés À, Collins MO, Reig-Viader R, Gou G, Goulding D, Izquierdo A, et al. Evolution of complexity in the zebrafish synapse proteome. Nat Commun. 2017 Mar 2;8(1):14613.

133. Georgijevic S, Subramanian Y, Rollins EL, Starovic-Subota O, Tang ACY, Childs SJ. Spatiotemporal expression of smooth muscle markers in developing zebrafish gut. Developmental Dynamics. 2007;236(6):1623–32.

134. Bolhassani A, Agi E. Heat shock proteins in infection. Clinica Chimica Acta. 2019 Nov;498:90– 100.

135. Okura GC, Bharadwaj AG, Waisman DM. Recent Advances in Molecular and Cellular Functions of S100A10. Biomolecules. 2023 Sep 26;13(10):1450.

136. Lou Y, Han M, Liu H, Niu Y, Liang Y, Guo J, et al. Essential roles of S100A10 in Toll-like receptor signaling and immunity to infection. Cell Mol Immunol. 2020 Oct;17(10):1053–62.

137. Song Y, Meng Z, Zhang S, Li N, Hu W, Li H. miR-4739/ITGA10/PI3K signaling regulates differentiation and apoptosis of osteoblast. Regenerative Therapy. 2022 Dec;21:342–50.

138. Klaus C, Liao H, Allendorf DH, Brown GC, Neumann H. Sialylation acts as a checkpoint for innate immune responses in the central nervous system. Glia. 2021 Jul;69(7):1619–36.

139. Ding Y, Cui K, Han S, Hao T, Liu Y, Lai W, et al. Lysophosphatidylcholine acyltransferase 3 (LPCAT3) mediates palmitate-induced inflammation in macrophages of large yellow croaker (Larimichthys crocea). Fish & Shellfish Immunology. 2022 Jul;126:12–20.

140. Greenblatt SM, Liu F, Nimer SD. Arginine methyltransferases in normal and malignant hematopoiesis. Experimental Hematology. 2016 Jun;44(6):435–41.

141. Zhu J, Liu X, Cai X, Ouyang G, Fan S, Wang J, et al. Zebrafish *prmt7* negatively regulates antiviral responses by suppressing the retinoic acid-inducible gene-I-like receptor signaling. FASEB j. 2020 Jan;34(1):988–1000.

142. Wang X, Xu W, Zhu C, Cheng Y, Qi J. PRMT7 Inhibits the Proliferation and Migration of Gastric Cancer Cells by Suppressing the PI3K/AKT Pathway via PTEN. J Cancer. 2023;14(15):2833–44.

143. Sinha S, Chatterjee SS, Biswas M, Nag A, Banerjee D, De R, et al. SWI/SNF subunit expression heterogeneity in human aplastic anemia stem/progenitors. Experimental Hematology. 2018 Jun;62:39–44.e2.

144. Liu W, Wang Z, Liu S, Zhang X, Cao X, Jiang M. RNF138 inhibits late inflammatory gene transcription through degradation of SMARCC1 of the SWI/SNF complex. Cell Reports. 2023 Feb;42(2):112097.

145. Xiao ZM, Lv DJ, Yu Y zhong, Wang C, Xie T, Wang T, et al. SMARCC1 Suppresses Tumor Progression by Inhibiting the PI3K/AKT Signaling Pathway in Prostate Cancer. Front Cell Dev Biol. 2021 Jun 25;9:678967.

146. Rodero MP, Tesser A, Bartok E, Rice GI, Della Mina E, Depp M, et al. Type I interferon-mediated autoinflammation due to DNase II deficiency. Nat Commun. 2017 Dec 19;8(1):2176.

147. Dao KHT, Rotelli MD, Petersen CL, Kaech S, Nelson WD, Yates JE, et al. FANCL ubiquitinates β-catenin and enhances its nuclear function. Blood. 2012 Jul 12;120(2):323–34.

148. Landelouci K, Sinha S, Pépin G. Type-I Interferon Signaling in Fanconi Anemia. Front Cell Infect Microbiol. 2022 Feb 7;12:820273.

149. Dao KHT, Rotelli MD, Brown BR, Yates JE, Rantala J, Tognon C, et al. The PI3K/Akt1 pathway enhances steady-state levels of FANCL. Heldin CH, editor. MBoC. 2013 Aug 15;24(16):2582– 92.

150. Leveau C, Gajardo T, El-Daher MT, Cagnard N, Fischer A, De Saint Basile G, et al. Ttc7a regulates hematopoietic stem cell functions while controlling the stress-induced response. Haematologica. 2020 Jan;105(1):59–70.

151. Gajardo T, Bernard M, Lô M, Turck E, Leveau C, El-Daher MT, et al. Actin dynamics regulation by TTC7A/PI4KIIIα limits DNA damage and cell death under confinement. Journal of Allergy and Clinical Immunology. 2023 Oct;152(4):949–60.

152. Estin ML, Thompson SB, Traxinger B, Fisher MH, Friedman RS, Jacobelli J. Ena/VASP proteins regulate activated T-cell trafficking by promoting diapedesis during transendothelial migration. Proc Natl Acad Sci USA [Internet]. 2017 Apr 4 [cited 2024 Apr 26];114(14). Available from: https://pnas.org/doi/full/10.1073/pnas.1701886114

153. Su W, Lin XT, Zhao S, Zheng XQ, Zhou YQ, Xiao LL, et al. Tripartite motif-containing protein 46 accelerates influenza A H7N9 virus infection by promoting K48-linked ubiquitination of TBK1. Virol J. 2022 Nov 3;19(1):176.

154. Cao F, Liu M, Zhang QZ, Hao R. PHACTR4 regulates proliferation, migration and invasion of human hepatocellular carcinoma by inhibiting IL-6/Stat3 pathway. Eur Rev Med Pharmacol Sci. 2016 Aug;20(16):3392–9.

155. Chu JY, Dransfield I, Rossi AG, Vermeren S. Non-canonical PI3K-Cdc42-Pak-Mek-Erk Signaling Promotes Immune-Complex-Induced Apoptosis in Human Neutrophils. Cell Reports. 2016 Oct;17(2):374–86.

156. Beemiller P, Zhang Y, Mohan S, Levinsohn E, Gaeta I, Hoppe AD, et al. A Cdc42 Activation Cycle Coordinated by PI 3-Kinase during Fc Receptor-mediated Phagocytosis. Wang YL, editor. MBoC. 2010 Feb;21(3):470–80.

157. Wu Q, Bai S, Su M, Zhang Y, Chen X, Yue T, et al. HIVEP3 inhibits fate decision of CD8+ invariant NKT cells after positive selection. Journal of Leukocyte Biology. 2023 Sep 27;114(4):335–46.

158. Wang Y, Jin H, Wang Y, Yao Y, Yang C, Meng J, et al. Sult2b1 deficiency exacerbates ischemic stroke by promoting pro-inflammatory macrophage polarization in mice. Theranostics. 2021;11(20):10074–90.

159. Hong W, Guo F, Yang M, Xu D, Zhuang Z, Niu B, et al. Hydroxysteroid sulfotransferase 2B1 affects gastric epithelial function and carcinogenesis induced by a carcinogenic agent. Lipids Health Dis. 2019 Dec;18(1):203.

160. Liu C, Li X. Identification of hub genes and establishment of a diagnostic model in tuberculosis infection. AMB Expr. 2024 Apr 13;14(1):36.

161. Liu J, Zhu H. TMEM106A inhibits cell proliferation, migration, and induces apoptosis of lung cancer cells. J of Cellular Biochemistry. 2019 May;120(5):7825–33.

162. Pong Ng H, Kim G, Ricky Chan E, Dunwoodie SL, Mahabeleshwar GH. CITED2 limits pathogenic inflammatory gene programs in myeloid cells. FASEB j. 2020 Sep;34(9):12100–13.

163. Yang J, Cao D, Zhang Y, Ou R, Yin Z, Liu Y, et al. The role of phosphorylation of MLF2 at serine 24 in BCR-ABL leukemogenesis. Cancer Gene Ther. 2020 Feb;27(1–2):98–107.

